# Gut transit and gut microbiome changes occur prior to the onset of motor impairment in a mouse model of Machado-Joseph disease

**DOI:** 10.1101/2025.04.30.651362

**Authors:** Vasilisa Zvyagina, Andrea Kuriakose, Ignacio Simo, Julia Y. Kam, Prapti Chakraborty, Katherine J. Robinson, Anastasiya Potapenko, Maxinne Watchon, Ian T. Paulsen, Hasinika K.A.H. Gamage, Angela S. Laird

**Author notes:** denote equal contributions (Vasilisa Zvyagina, Andrea Kuriakose). denote equal contributions (Hasinika K.A.H. Gamage, Angela S. Laird).

## Abstract

We previously identified microbial shifts prior to the onset of motor and neurological symptoms within a mouse model of the fatal neurodegenerative disease Machado-Joseph disease (MJD). Here, we aimed to explore possible mechanisms contributing to these changes within the microbiome-gut-brain axis, and whether it preceded or followed central neurodegeneration. Here, we report that pre-symptomatic male MJD mice present with significantly different microbiome communities as early as 5-weeks-old. Furthermore, we show that male MJD mice have faster total gut transit times by 9-weeks-old, prior to signs of impaired motor function by 11-weeks-old. To elucidate whether these microbial and colonic functional changes are due to the presence of pathological and morphological changes in the gut, we quantified the formation of ataxin-3 protein aggregates within the gut and examined morphological changes within the gut of pre- and early symptomatic MJD mice relative to proteinopathy in the brain. Interestingly, we observed ataxin-3 aggregates within the brains of pre-symptomatic MJD mice, with significantly more aggregates present in MJD than WT mice from 7-weeks-of-age, an earlier timepoint than previously reported, coinciding with changes within the microbiome. However, we observed no ataxin-3 protein aggregates and no changes in enteric neuron populations or morphology within the gut. Analysis of endocrine factors involved in gut motility and inflammatory markers within the small intestine of 13-week-old males revealed increased expression of genes encoding cholecystokinin (*Cck*), ghrelin (*Ghrl*), heme oxygenase (*Ho1*), interleukin-1 beta (*Il1b*), and decreased inducible nitric oxide synthase (*Nos2*). Together, we demonstrate for the first time that colonic dysfunction occurs after gut microbiome changes, but prior to the onset of motor impairments in male MJD mice. Our work suggests that whilst proteinopathy or morphological changes within the gut may not be involved in these changes, inflammation and related endocrine changes could have a role in the interplay between the gut and brain during MJD development, warranting further investigation.

## Introduction

A growing body of evidence indicates that alterations to the gut microbiome contribute to the pathogenesis and pathophysiology of neurodegenerative diseases [1–9]. This relationship is hypothesised to involve various mechanisms, including bi-directional communication between the gut and brain [10]. This interplay between the microbiome-gut-brain axis is evident in neurodegenerative diseases such as Parkinson’s disease (PD) and Huntington’s disease (HD), where alterations in the gut microbiome have been implicated in the pathogenesis and progression of disease [11–13]. Findings from clinical studies suggest that gastrointestinal symptoms such as bowel dysfunction are present in both PD and HD patients, with recent studies highlighting the presence of gastrointestinal dysfunction years prior to the onset of motor symptoms [14].

We have recently studied the composition of the gut microbiome of mice modelling the fatal neurodegenerative disease Machado Joseph disease (MJD), also known as spinocerebellar ataxia type-3 (SCA3) [2]. MJD is characterised by motor symptoms such as ataxia, rigidity, dystonia and ophthalmoplegia [15, 16]. The cause of MJD is known to be the inheritance of an abnormal form of the *ATXN3* gene carrying an expanded trinucleotide (CAG) repeat region [17]. In healthy individuals, the *ATXN3* gene contains approximately 12 to 43 CAG repeats, while greater than 60 repeats have been reported in MJD patients [18, 19]. Through examining the gut microbiome in MJD, we have reported that male and female transgenic MJD mice exhibit significant shifts in their gut bacterial community. Notably, these microbiome alterations were observed at pre-symptomatic stages of the disease, prior to the onset of known neurological and motor symptoms [2]. Furthermore, we found strong correlations between pre-symptomatic gut microbiome shifts and the severity of neurological symptoms present at later stages of MJD progression [2].

As MJD is a genetic condition, it is not yet understood what causes these microbiome changes to occur. Possible triggers may include gut motility changes, potentially resulting from neurodegeneration within enteric neurons, or morphological changes within the small intestine and colon. Changes in gut motility, such as constipation, have previously been reported in early Parkinson’s disease patients [1], and have been proposed to trigger gut microbiome alterations [20]. Other potential causes may include altered gut membrane integrity, changes to inflammatory signalling or neurotransmitter abundance within the gut, which may impact the microbiome directly, or indirectly, consequently impacting gut motility. Disruptions in this gut– brain axis can lead to altered immune responses, including the activation of inflammatory pathways that may exacerbate neurodegenerative processes [21]. Similarly, while this has not been directly tested in neurodegenerative disease, gut microbiome changes are proposed to occur within inflammatory bowel disease and anxiety disorders, due to alterations in pro-inflammatory cytokines and neurotransmitters, respectively [21–23].

Within this study, we aimed to examine the potential triggers of the altered faecal microbiome in MJD mice and identify whether these occur prior to central neurodegeneration. Hence, we examined central neurodegeneration, gut function (motility and faecal output), microbiome composition, gut morphology and signs of enteric neurodegeneration at a range of disease stages to increase the understanding of these potential sequalae in the context of MJD. Here, we report for the first time that gut microbiome and colonic functional changes precedes motor impairment in MJD, with a potential link with inflammation and associated endocrine changes. The findings of this study enhance whether changes in the gut microbiome have an impact on disease progression and neurodegeneration in MJD, and whether further investigation into methods to ameliorate these changes are warranted.

## METHODS

### Experimental animals

This study utilised CMVMJD135 transgenic mice that express human *ATXN3* (h*ATXN3*) with an expanded CAG repeat sequence, driven by a CMV promotor at near endogenous levels [24]. The animal procedures performed within this study were approved by the Macquarie University Animal Ethics Committee (ARA 2022/008). Mouse tail tissue was collected at approximately 2-3 weeks-of-age and DNA was extracted to confirm presence or absence of the CMV promoter. DNA from animals identified to be CMV-positive underwent further analysis to sequence the CAG repeat region of the h*ATXN3* gene. Animals expressing h*ATXN3* with CAG lengths ranging from 133 to 149 were recruited into the study. CAG repeats of all mice recruited into the study was statistically analysed (data not shown) to confirm that variation of CAG repeats was not statistically significant across sub-cohorts.

The CMVMJD135 mouse colony was maintained at Australian BioResources (Moss Vale, Australia) and animals were shipped to Macquarie University from 4 weeks-of-age for commencement of experimentation at 5 weeks-of-age. A total of 95 MJD mice were recruited into the study. This included two cohorts of mice to achieve different experimental aims. The first cohort of mice (male MJD n = 11, male WT n = 12, female MJD n = 12, female WT n = 12) was used to examine how motor, colonic function and the gut microbiome change through presymptomatic to early stages of MJD, with euthanasia at 13-weeks-of-age and sample collection of brain, colon and small intestine for immunohistochemistry and RT-qPCR (Figure 1A). A second cohort of male mice was euthanised at 5, 7, 9 or 11 weeks-of-age (MJD n = 6 and WT n = 6 per timepoint) to allow collection of tissue from a range of ages for immunohistochemical, protein and RNA samples from the brain, colon and small intestine (Figure 4A). All mice were group-housed (2–5 mice per cage) with access to environmental enrichment and standard chow diet throughout the duration of the study. Food and water were available *ad libitum*. To reduce any confounding variables on the gut microbiome due to coprophagy, mice were housed relative to sex and genotype.

**Fig. 1.**
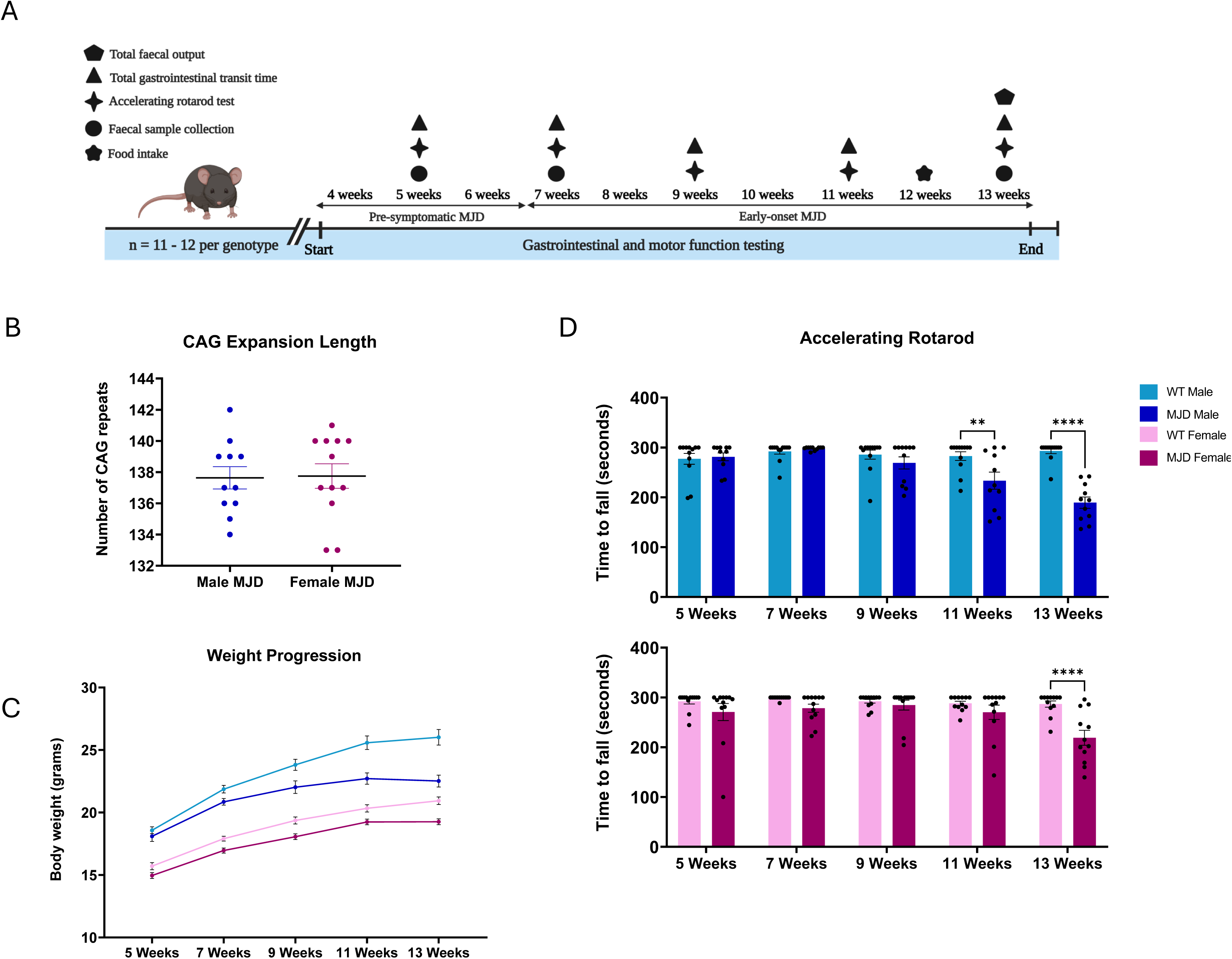
Transgenic male and female MJD mouse models expressing CAG-repeat expanded human *ATXN3* develop motor function impairment that progressively increases with age. **(A)** Timeline of the experiment. A total of 47 mice (n =12/group in male wildtype (WT), female WT and female MJD groups, and n = 11 in male MJD group) were used to collect the samples and data at the indicated timepoints. Timeline figure was created using BioRender.com. **(B)** CAG-repeat expansion length comparison between male and female MJD mice showed no significant differences. **(C)** Male MJD group showed significantly lower body weight progression compared to the male WT mice from 11 weeks-of-age, whereas no significant differences were found between female MJD and female WT mice. **(D)** Male MJD and male WT mice demonstrated significant motor dysfunction during the accelerating rotarod test from 11 weeks-of-age. Female MJD mice developed motor impairment at a later age, where statistically significant differences between genotypes were observed for the accelerating rotarod test at 13 weeks-of-age. Data are presented as group mean ± standard error mean (SEM), ** p < 0.01 and **** p < 0.0001.

Animals were euthanised via sodium pentobarbitone overdose (300 mg/kg, IP) at either 13 weeks-of-age for Cohort 1 or at 5, 7, 9 or 11 weeks-of-age for Cohort 2 following all motor and gastrointestinal function testing. Intracardiac perfusion with 0.9 % (v/v) saline was performed, followed by extraction of the brain, small intestine, colon, and colonic faecal contents. Proximal and distal ends of each small intestine subregion and colon samples were collected and snap frozen in liquid nitrogen for protein and RNA extraction. The remaining length of small intestine and colon samples were post-fixed in 10 % neutral buffered formalin and whole brains in 4 % paraformaldehyde for immunohistochemical processing.

### Neurological impairment and motor behaviour dysfunction

All mice were monitored weekly for neurological and behavioural changes to ensure that appropriate animal welfare was maintained. For each animal within Cohort 1, motor coordination and stamina were assessed by the time taken to fall from an accelerating rotarod machine (Model 7650, Ugo Basile). The test began at 4 rpm and the speed of the accelerating rotarod was gradually increased to 40 rpm over a span of 300 s. The total time taken for a mouse to fall from the machine was recorded, with a maximum time of 300 s recorded if the mouse did not fall. Each mouse performed three trials fortnightly and the average of the two best times of each mouse was statistically analysed. This test was performed during the light cycle, between the hours of 8 am to 2 pm, with minimum 5 mins of rest between each trial to prevent fatigue.

### Total gastrointestinal transit time

Total gastrointestinal transit was examined for each mouse in Cohort 1 by determining the amount of time taken to detect Carmine Red, a dye that cannot be absorbed by the lumen of the gut, in faeces following oral administration. 6 % (w/v) Carmine solution (Carmine red dye dissolved in 0.5 % methylcellulose, Sigma Aldrich) was administered at a volume of 150 μL to each mouse via oral gavage between 9 am to 12 pm. Following administration, each mouse was individually housed in clean cages with food and water provided *ad libitum*. Mice were closely monitored for red faecal pellets and confirmation smears were conducted on white paper towels. Total gut transit time was calculated as the time interval between administration and passage of the first red or partially red faecal pellet.

### Food intake measurements

To eliminate food intake as a confounding variable affecting gastrointestinal function tests, the average food intake of each cage was measured from a subset of Cohort 1 mice. A total of n = 12 male (6 MJD and 6 WT controls) and n = 11 female (4 MJD and 7 WT controls) 12-week-old mice were used to measure feed intake from each cage over a 24-hr period.

### Total faecal output

Total faecal output was assessed for each mouse in Cohort 1 by counting the number of faecal pellets produced in one hour (with food and water provided *ad libitum)*. The wet and dry weight of the faecal output, and fluid content of the total faecal output was calculated. Faecal pellets were collected every 10 min to avoid excess fluid absorption (e.g., from urine) for faecal pellet fluid content measurements. Faecal pellets were placed into a 65 °C oven overnight and dry weight was determined. Fluid content was calculated using the following equation: % fluid content = [(wet weight – dry weight)/ wet weight] × 100 [25].

### 16S rRNA amplicon sequencing of the gut microbiome

Faecal samples were collected from each mouse under aseptic conditions at weeks 5, 7, and 13. Total DNA was extracted from faecal samples using the DNeasy PowerLyzer Powersoil kit (Qiagen, Germany) according to the manufacturer’s instructions. The V4 region of the 16S rRNA gene was amplified using 515F (5′-GTGCCAGCMGCCGCGGTAA-3′) and 806R (5′-GGACTACHVGGGTWTCTAAT-3′) primers with Golay barcodes and the Platinum^TM^ Hot Start PCR master mix (Thermo Fisher Scientific, Australia). Amplicon concentrations were measured using the Quant-iT^TM^ PicoGreen® dsDNA assay kit (Invitrogen, Australia), equimolar volumes of barcoded amplicons from each sample were then pooled, and amplicons were gel purified using the Wizard® SV gel and PCR cleanup system (Promega, Australia). Multiplexed amplicons were sequenced on the Illumina Miseq (2 x 250 bp) platform at the Ramaciotti Centre for Genomics, Australia.

### Bioinformatic analyses of microbiome sequence data

Raw sequencing data was processed using Quantitative Insights Into Microbial Ecology software (QIIME2 version 2022.2) [26]. De-multiplexed reads were quality-filtered (median of quality score > 30 and read length > 154 bp), and denoised using Deblur [27]. The identified bacterial amplicon sequence variants (ASVs) were aligned using Multiple Alignment using Fast Fourier Transform (MAFFT) and a phylogenetic tree was constructed using FastTree2 [28, 29]. The SILVA 138 classifier of the 515F/806R region of the 16S rRNA gene was used to assign taxonomy with 99% similarity [30]. Following quality filtering, a total of 9,326,373 reads were retained, and each sample was rarefied at 20,000 reads prior to statistical analysis.

To present the ordination of the overall gut microbiome community structure, non-multidimensional scaling (nMDS) plots were used. The vegan R package [31] was used to obtain nMDS ordinates based on Bray-Curtis similarity matrices of the abundance of ASVs. To examine statistically significant shifts in the gut microbiota community structure across sex, genotype, and weeks of age, permutational multivariate analysis of variance (PERMANOVA) tests with 9999 permutations were used. To assess the alpha diversity of the faecal microbiome, Shannon’s diversity index, Simpson’s evenness index, and Faith PD’s richness index were calculated using QIIME2. Plots were constructed using GraphPad Prism (version 9.5.1) software (GraphPad Software, La Jolla California, USA).

To identify ASVs with significantly different (*p* < 0.05) abundances between genotypes at 5, 7, and 13 weeks, Linear Discriminant Analysis effect size (LEfSe) (online Galaxy Version 1.0) was used [32]. Male and female mice were analysed separately. MJD and WT were used as classes with no subclasses or subjects. A Linear Discriminant Analysis score was identified for each ASV using the following default parameters: *p* < 0.05 for the factorial Kruskal-Wallis test among classes, *p* < 0.05 for the pairwise Wilcoxon test between subclasses, and < 2.0 threshold for the logarithmic LDA score for discriminative features. Multiple comparison adjustment was performed for all p-values using Benjamini and Hochberg’s false discovery rate (5%) correction. The abundance (Log10 transformed) of ASVs that were found to be significantly different between groups were used to construct unpaired heatmaps using GraphPad Prism software.

### Correlation network analyses

Pairwise correlation analyses were used to examine the associations between early-stage gut microbiome changes with the onset of behavioural and gut functional impairment. For this, the relative abundances of ASVs that demonstrated statistically significant diuerences between MJD and WT groups at 5-weeks-of-age were used to examine their correlations with gut transit measurements at 9-weeks-of-age and rotarod data collected at 11 weeks-of-age using the Hmisc R package [33]. Statistically significant correlations (*p* > 0.05, FDR 5 %) were used to construct correlation networks using Cytoscape software (version 3.10.1).

### Confirmation of ataxin-3 expression in CMVMJD135 mice

The expression of expanded human ataxin-3 that is driven by the ubiquitous CMV promotor in the CMVMJD135 mice used within this study has been previously confirmed in the CNS but not in other organs such as the small intestine and colon [24]. To confirm presence of h*ATXN3* in transgenic animals within the small intestine and colon, we performed RT-PCR on samples from 13-week-old male mice (n = 1 WT and n = 1 MJD) and western blotting for human ataxin-3 in the small intestine (n = 7 WT and n = 8 MJD). Primers for human *ATXN3* (Table 1) were designed using primer3web version 4.1.0 (https://primer3.ut.ee) based on sequences obtained from the Human Genome (NCBI). After euthanasia, small intestine and colon tissue (100 mg) was immediately homogenized in 1 mL of TRIzol REAGENT (Invitrogen) for total RNA extraction following the manufacturer’s instructions. RNA quantification and purity were determined using NanoDrop 2000 UV-Vis spectrophotometer (Thermofisher Scientific). To eliminate possible genomic DNA contamination, all samples were treated with DNase I (Promega, Madison, WI, USA) starting from 5 µg of total RNA according to the manufacturer’s instructions. Then, first-strand cDNA synthesis was performed using Oligo dT as the primer with Superscript III (Thermofisher Scientific) for 60 min at 50 °C followed by 15 min at 85 °C again following the manufacturer’s instructions and finally diluted to a final volume of 150 µL. RT-PCR reactions were conducted on an Eppendorf Mastercycler nexus in a final volume of 20 µL using Taq polymerase (Bioline), 0.5 µM each primer, 2 µL of cDNA template, dNTPs 0.25 mM each and 1.5 mM Mg^2+^. PCR products were analysed by electrophoresis in 2 % agarose gel stained with SYBR Safe (Thermofisher Scientific) and digitalized in a Gel Doc™ EZ System (BioRad).

**Table 1.**
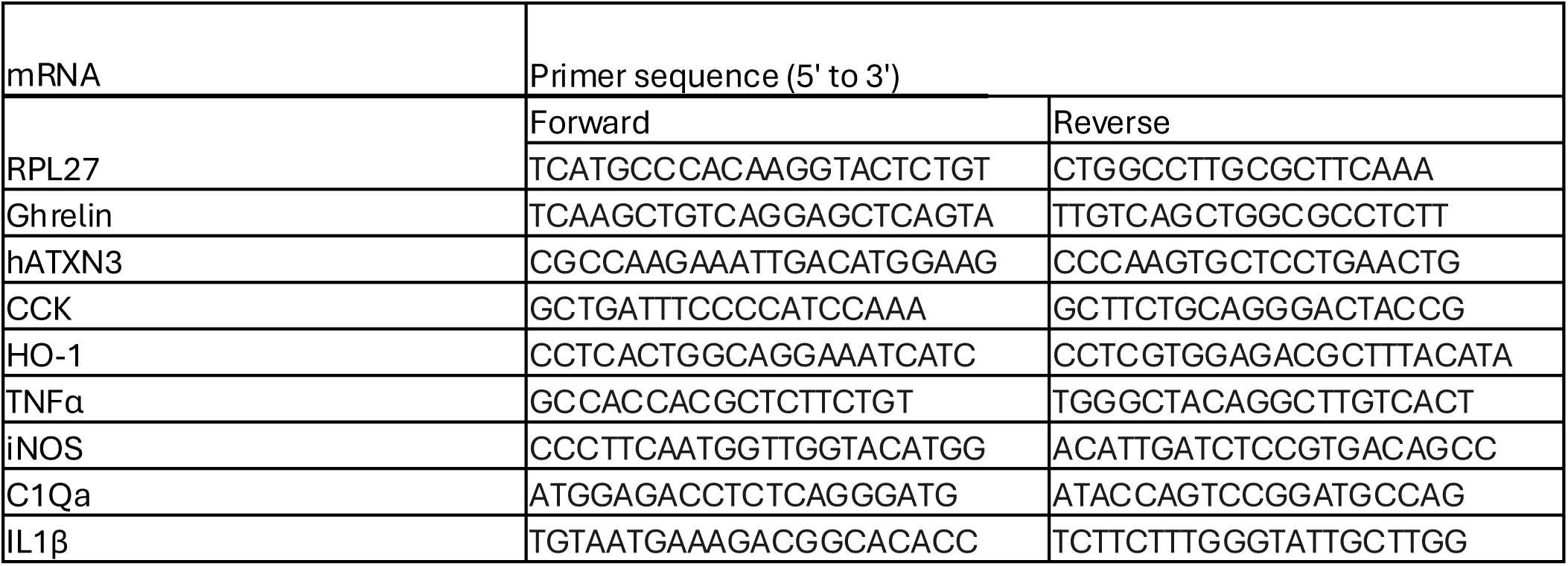
Primer sequences used for end point PCR and for real-time quantitative reverse transcriptase polymerase chain reaction (RT-qPCR)

To confirm the expression of human ataxin-3 in the small intestine, snap frozen tissue collected from Cohort 1 mice at euthanasia was placed in 20 µL/mg of RIPA containing cOmplete Protease Inhibitor Cocktail (Roche) and PhosSTOP Phosphatase Inhibitor Cocktail tablets (Roche). The small intestine was first broken into small pieces using surgical micro scissors, then probe sonicated using an Omniruptor 250 Ultrasonic Homogeniser (Omni International), centrifuged at 13,200 rpm at 4 °C for 25 min, and the supernatant containing RIPA-soluble protein was collected. The total protein concentration was determined using a Pierce BCA protein assay kit (ThermoFisher Scientific) according to the manufacturer’s instructions.

10 µg of protein was prepared with NuPage Sample Reducing Agent (Invitrogen) containing dithiothreitol and Laemmli Sample Buffer (Bio-Rad), denatured and separated using a 4-12 % acrylamide gradient NuPAGE Bis-Tris gel (Invitrogen). Separated proteins were transferred to a 0.45 µm polyvinylidene difluoride (PVDF; Cytiva) membrane, followed by ataxin-3 labelling using a rabbit anti-ataxin-3 antibody (gifted by H. Paulson). The immunoblots were then incubated in goat anti-rabbit IgG (H+L) HRP Conjugate (Promega, #W4011). Protein bands were visualised on ImageQuant LAS 4000 (Cytiva) by chemiluminescent reaction with Clarity Western ECL Substrate (Bio-Rad). Digital 8-bit TIFF images were exported to ImageStudio (LI-COR Biosciences) for full length human ataxin-3 quantification and normalised against a GAPDH loading control (mouse anti-GAPDH antibody (#60004-1-Ig, Proteintech) with goat anti-mouse IgG (H+L) HRP Conjugate (Promega, #W4021) secondary).

### Immunohistochemical staining

Following euthanasia at 13-weeks-of-age of male Cohort 1 animals (MJD n = 6 and WT n = 6) and Cohort 2 animals (MJD n = 6 and WT n = 6 per time point), tissue samples from the brain, small intestine and colon were extracted. Brains were preserved in 4 % paraformaldehyde (PFA) and small intestine and colon samples preserved in 10 % neutral buffered formalin. Following 24 hrs in 4 % PFA, brains were briefly washed in PBS (3 × 2 min) then placed in 30 % sucrose solution for a further 24 hrs. Brains were then embedded in Tissue Tek OCT compound (Sakura Finetek) and cryosectioned at 35 µm before storage in PBS at 4 °C until use. Following 24 hrs in 10 % neutral buffered formalin, small intestine and colon samples were briefly washed in PBS (3 × 2 min) then processed overnight in an automated tissue processor prior to embedding in paraffin using the “bundling” technique as described by Williams *et al.* [34]. Sections were sliced on the microtome at 10 µm and collected slices underwent deparaffinization prior to staining for hematoxylin and eosin (H&E) stain, or for immunohistochemistry as described below.

Collected cryosections and formalin fixed paraffin embedded samples underwent 3, 3’ diaminobenzidine (DAB) immunohistochemistry for ataxin-3 or HuC immunoreactivity. Samples were first incubated in 50 % ethanol for 20 min at RT, then quenched in 3 % H2O2 + 50 % ethanol for 30 min at 4 °C prior to blocking in 3 % bovine serum albumin (BSA) + 0.25 % or 0.5 % Triton X-100 in PBS for 1hr at RT. Sections were incubated in either rabbit anti-ataxin-3 (1:2000 for brains and 1:1000 for intestine samples, gift from H. Paulson) for 24 hrs at 4 °C or for 72 hrs at 4 °C in mouse anti-HuC (1:500, Invitrogen, Cat#A-21271) diluted in blocking solution, followed by overnight incubation at 4 °C in biotinylated secondary antibody (1:2000, Vector Laboratories; goat anti-rabbit BA-1000 or goat anti-mouse BA-9200) diluted in blocking solution. Sections were then incubated for 1 hr at RT in Avidin-Biotin complex (Vector Laboratories, Vectastain Elite ABC Kit, Peroxidase Cat#PK-6100), prepared as per manufacturer instructions and left to complex for 30 min before application. Ataxin-3 and HuC immunolabelling was detected by incubating in DAB chromogen (Vector Laboratories, DAB Substrate Kit, Peroxidase Cat#SK-4100) for 5 min. In preparation for imaging, cryosections were mounted onto glass slides then cover slipped with DAKO mounting media, and formalin fixed paraffin embedded slides cover slipped with DPX mounting media.

### Microscopic imaging and analysis

Whole slide imaging of brain and colon ataxin-3 DAB sections, HuC DAB, H&E small intestine sections and Sudan III-stained faecal samples were performed under brightfield (Olympus VS200 MTL, UPLXAPO 20x objective lens running VS200 ASW 4.1.1 software) at optimised exposure times.

Quantification of ataxin-3 aggregates within the medulla oblongata was performed by a blinded investigator using the Multi-Point counting tool on ImageJ. Required regions of interest were identified using the Allen Brain Atlas [35]. Three anatomically similar sections were chosen per animal for manual aggregate quantification within the medulla oblongata. The sum of the aggregates from the three sections was taken as the total number of aggregates for the animal.

Colon and small intestine muscle layer thickness was analysed using the Trainable WEKA Segmentation plugin on FIJI [36], which was used to select the muscularis layer within each section of the H&E stained bundle. The ImageJ Local Thickness tool was used to measure the thickness of the muscularis layer across the whole image, and the mean value was obtained from the Histogram tool output for statistical analysis. The number of enteric neurons within a small intestine bundle cross-section, per animal, was quantified using an ImageJ protocol involving the “Analyse Particles” function.

### Reverse Transcriptase Quantitative PCR (RT-qPCR)

Samples were suspended in TRIzol™ Reagent (Invitrogen, cat 15596026) and finely minced using a sharp lancet. Tissue was then disrupted and homogenized using a Tissue Grinder Potter-Elvehjem. The samples were centrifuged at 13,000 rpm for 15 min, and the supernatant transferred to a new tube. Total RNA was isolated following the manufacturer’s protocol for TRIzol reagent. RNA samples were treated with DNase I (RQ1 DNase, Promega M6101) for 30 min to remove DNA contamination. RNA purity and concentration were assessed using a NanoDrop spectrophotometer (Thermo Fisher Scientific) and a 260/280 ratio of 2.0 was considered acceptable. Reverse transcription of 5 µg of total RNA to cDNA was performed using the High-Capacity cDNA Reverse Transcription Kit (Applied Biosystems, Thermo Fisher Scientific, 4368814) in a total reaction volume of 20 µL. The cDNA was diluted to a final volume of 150 µL. Primers were designed using primer3web version 4.1.0 (primer://primer3.ut.ee) based on the sequences obtained from NCBI. Prior to real-time RT-qPCR analysis, the efficiency of amplification for all primer pairs was evaluated to ensure proper use of the ΔΔCT method. RT-qPCR reactions were performed on QuantStudio7 Pro (Applied Biosystems) in a 10 µL final volume comprised of 1 µL of cDNA template, 5 µL of 2X SYBR Green master mix (ThermoFisher Scientific, cat# 4367659), 0.5 µL of 10 µM forward and reverse primer mix and sterile water. Melting curve analysis was conducted to confirm the specificity of the real-time RT-qPCR reaction, which was programmed to include a melting profile immediately following the thermal cycling protocol. Mean CT values and standard deviations were used for ΔΔCT calculations. Fold differences in gene expression were calculated using the ΔΔCT method [37]. Gene expression was normalized to the endogenous internal control gene Ribosomal Protein L27 (*Rpl27*). The results are presented as the fold change of target gene expression relative to a reference sample and normalized to the reference gene.

For RT-qPCR analysis of *Ghrl* and *Cck*, RNA was extracted from the small intestine of n = 5 WT and n = 5 MJD mice from Cohort 1. However, one RNA extraction did not yield sufficient quality for analysis, resulting in a final sample size of n = 4 WT and n = 5 MJD 13-week-old mice. For RT-qPCR analysis of inflammatory markers, a new set of RNA extractions was performed using n = 5 WT and n = 5 MJD 13-week-old mice from Cohort 1 resulting in a final sample size of n = 5 per group.

### Statistical Analyses

Data analysis was performed using GraphPad Prism (version 10). Body weight progression and total neurological score were analysed using a mixed-effects model with factors of sex, genotype, and a repeated measure of age. Post hoc analyses were conducted using Tukey’s multiple comparisons test. Total gastrointestinal transit time, accelerating rotarod, alpha diversity (Shannon’s diversity, Faith’s PD richness, and Simpson’s evenness indices), and the ratio between the relative abundance of *Firmicutes* and *Bacteroidota* phyla were analysed using two-way analysis of variance (ANOVA), with the factors being genotype and age, across varying timepoints during MJD progression. Post-hoc analyses were performed using Šídák’s multiple comparisons test. Statistical significance of the total faecal output (total pellet counts, total wet and dry weight of faecal pellets, and total fluid content of faecal pellets) was analysed using an ordinary one-way ANOVA. Post-hoc analyses were conducted using Tukey’s multiple comparisons test. The number of inherited CAG repeats in male and female MJD mice was compared using an unpaired t-test. The amount of full-length human ataxin-3 protein in the brain and small intestine of male MJD mice was compared to male WT littermates using an unpaired t-test. Statistical significance of the total number of ataxin-3 aggregates within the medulla oblongata of male MJD mice was compared to male WT littermates using a two-way ANOVA, followed by Tukey’s post-hoc analysis. Differences in small intestine muscularis layer thickness and total HuC+ neurons in the small intestine between WT and MJD mice were analysed using an unpaired t-test. Colon muscularis layer thickness in WT and MJD mice across different timepoints was analysed using a two-way ANOVA, followed by Tukey’s post-hoc analysis. All RT-qPCRs were analysed using an unpaired t-test.

## RESULTS

### Male and female MJD mice develop motor impairment at different ages

Genetic sequencing of Cohort 1 mice to confirm CAG repeat length revealed that the MJD animals carried repeat lengths varying from 133 to 142. We did not detect a statistically significant difference when comparing the CAG repeat length within h*ATXN3* across male and female MJD mice (p = 0.9617, **Figure 1B**), suggesting any sex dependent effects are not due to underlying differences in CAG repeat length inheritance. Statistical comparison of body weight progression demonstrated a significant effect of genotype (p < 0.0001), sex (p < 0.0001) and age (p < 0.0001). Statistically significant age x genotype and genotype x sex interaction effects were also detected, (p = 0.0010 and p = 0.0163, respectively). Male WT mice gained weight at a consistent rate from 5 to 13 weeks of age (**Fig. 1C**), whereas male MJD mice either remained at a stable weight or began to lose weight from 11 weeks-of-age. Body weight was found to be significantly lower in male MJD mice when compared to male WT mice at 11- and 13-weeks-old (p < 0.0001 and (p < 0.0001, respectively). While body weight was found to be significantly lower in female mice when compared to male mice, (p < 0.0001), no statistically significant differences were identified between female MJD and WT groups at any of the examined time points.

Motor coordination, endurance and stamina of MJD mice compared to WT littermates was assessed on the accelerating rotarod. Due to sex-specific differences observed in body weight progression in this study and previous observations on sex-dependent differences in motor performance and progression [2], the motor function of male and female mice was analysed separately. Significant effects of genotype (p < 0.0001) and age (p < 0.0001) were observed on rotarod performance, as well as a significant interaction between genotype and age (p < 0.0001), whereby male MJD mice fell off the rotarod at a faster rate as the disease progressed. Post-hoc comparisons confirmed this where male MJD mice showed a significantly shorter time spent on the rotarod when compared to WT males at 11 weeks of age (p = 0.0025) and 13 weeks-of-age (p < 0.0001, **Fig. 1D**). Analysis of rotarod performance in female mice also revealed significant main effects of genotype (p < 0.0001) and age (p = 0.0025), and a significant genotype and age interaction effect (p = 0.0306). Post-hoc comparisons revealed a statistically significant difference in performance at 13 weeks-of-age, whereby female MJD mice showed significant reductions in the time taken to fall off the rotarod only at 13 weeks-of-age (p < 0.0001, **Fig. 1D**) compared to female WT controls. Overall, female MJD mice demonstrated motor dysfunction during rotarod performance at a later stage of disease compared to male MJD mice, who displayed motor dysfunction from 11-weeks-of-age.

### Male MJD mice develop altered gut function at early disease stages

We examined gut function of male and female MJD mice compared to non-transgenic WT littermate controls using two measures of gut function: total gut transit time and total faecal output. First, we examined the time taken for food to pass through the gastrointestinal tract. A significant effect of genotype (p < 0.0001) and an interaction between genotype and age (p < 0.0001) was seen, where male MJD mice had a significantly faster transit of food through the gastrointestinal tract as they aged. A statistically significant effect of age was not detected in male mice (p = 0.5789). Total gut transit was similar in female MJD mice and female WT mice, with no statistically significant differences detected (p = 0.1547) in the total gut transit compared to female WT mice. Post-hoc comparisons revealed that male MJD mice displayed significantly faster gastrointestinal transit at 9 weeks-of-age (p = 0.0025), 11 weeks-of-age (p = 0.0150), and 13 weeks-of-age (p < 0.0001) compared to male WT mice (**Fig. 2A**).

**Fig. 2.**
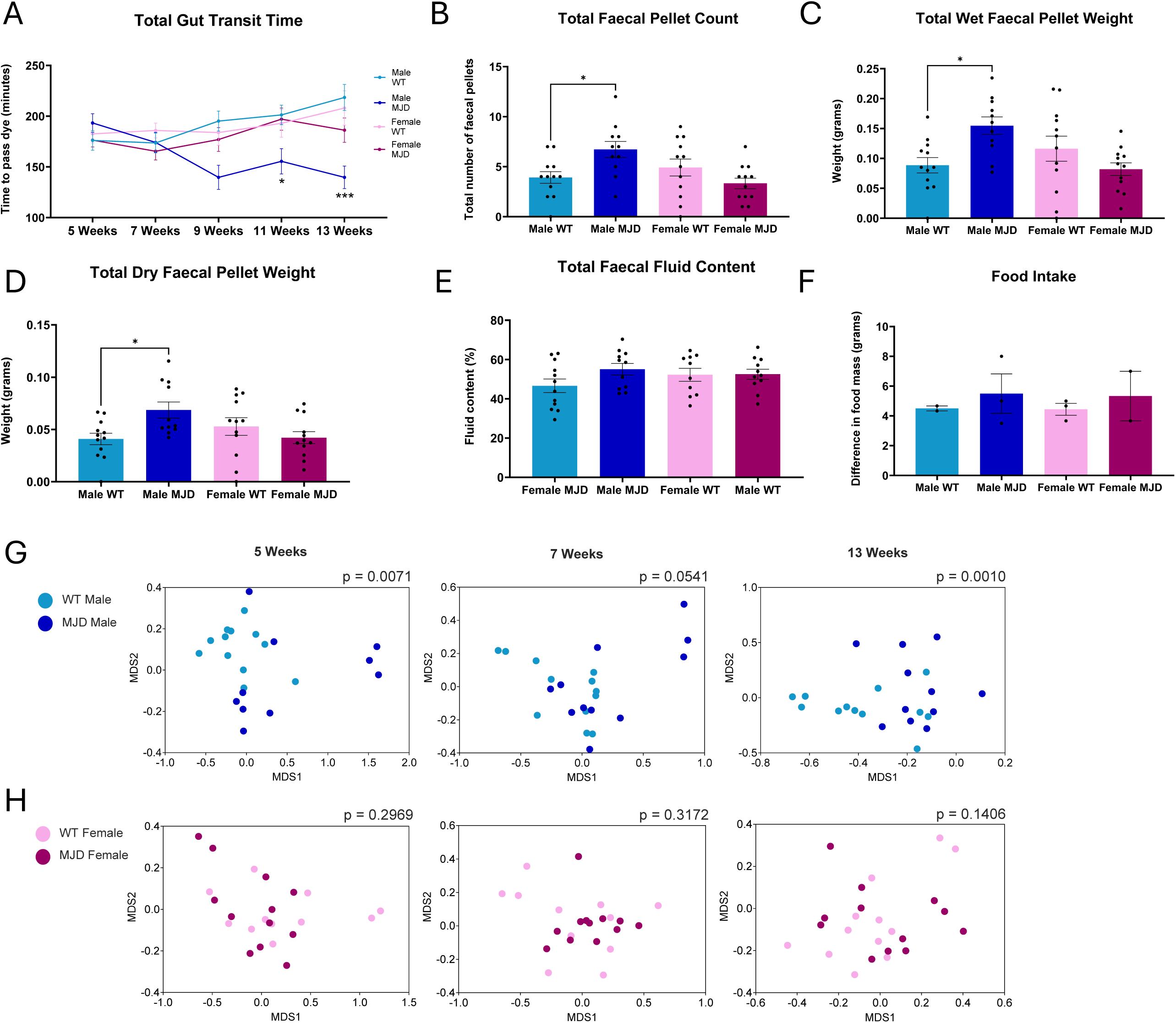
Male MJD mice develop colonic dysfunction and gut microbiome changes that increase with disease progression. **(A)** Male MJD mice demonstrate significantly faster total gut transit (transit of food through the gastrointestinal tract) compared to WT male controls. Male MJD mice also demonstrate a statistically significant increase in **(B)** total faecal pellet count, and **(C)** total wet weight and **(D)** total dry weight of faecal pellets compared to male WT controls. Neither sex showed statistically significant differences in the **(E)** total fluid content of faeces nor **(F)** food intake measurements between MJD and WT groups. Female MJD and female WT mice did not display any significant differences for any of the gut function tests above. Data are presented as group mean ± SEM, n = 11-12 per group, except for **(G)** where a subset of mice was used: n = 12 male (6 MJD and 6 WT controls) and n = 11 female (4 MJD and 7 WT controls) and (F) where cages were weighed (n = 2 - 3 cages per group). p < 0.05, **p < 0.01, and **** p < 0.0001. Ordinates of the gut microbiota community structure are visualised using Bray-Curtis similarity-based non-multidimensional scaling (nMDS) plots. The gut microbiota community structure of **(G)** male mice showed significant differences between MJD and WT starting from week 5, whereas **(H)** female mice showed no significant differences between the MJD and WT groups at 5, 7 and 13-weeks-of-age. *p* values were obtained using PERMANOVA analysis between groups.

At 13 weeks-of-age, faecal output of each individual mouse was calculated and used to examine the total faecal pellet count, total wet and dry faecal pellet weight, and total faecal fluid content. The number of faecal pellets produced within 1 hr revealed a significant difference across groups (p = 0.0088). Post hoc-analysis revealed that in comparison to male WT controls, male MJD mice had a significantly greater (p = 0.0359, **Fig. 2B**) total output of faecal pellets. Analysis of the total wet weight of faecal pellets also revealed a significant difference among groups (p = 0.0076). Post-hoc comparisons revealed increased faecal pellet weight in male MJD mice when compared to WT control mice (p = 0.0212, **Fig. 2C**). The dry weight of the faecal pellets also showed a significant difference between groups (p = 0.0289). Comparison of total dry faecal pellet weights of male MJD mice and male WT mice showed a significantly greater dry mass produced by male MJD mice compared to the control group (p = 0.0378, **Fig. 2D**). Analysis of faecal fluid content revealed no significant differences between male MJD and male WT controls (**Fig. 2E**). Interestingly, female MJD mice had no significant differences in any of the tested parameters of faecal output compared to female WT mice.

To validate gastrointestinal function testing and eliminate the amount of food intake as a confounding variable, food intake measurements were taken from a subset of mice at 12 weeks-of-age. Upon examination of food intake, no statistically significant differences between male MJD and male WT mice, nor between female MJD and female WT mice were found (**Fig. 2F**). Normalising food intake relative to the mean body weight of each cage also did not result in any significant differences.

### Male MJD mice display shifts in overall gut microbiome community structure

Faecal samples collected at 5, 7 and 13 weeks-of-age were used to profile the gut microbiome using 16S rRNA amplicon sequencing. The microbiome beta diversity of male MJD mice was significantly different compared to that of male WT controls (**Fig. 2G**), demonstrating shifts in the microbiome community structure with MJD development These shifts in the gut microbial community structure were seen at 5 weeks (p = 0.0071, PERMANOVA) and 7 weeks (p = 0.0541), and became more profound by 13 weeks of age (p = 0.0010). In contrast, female MJD mice and female WT mice did not demonstrate any significant differences in beta diversity at any of the three tested time points (**Fig. 2H**).

We next examined the alpha diversity of the gut microbiome community using Shannon’s diversity index (**Supp Fig. 1A**)., Faith’s PD richness index (**Supp Fig. 1B**). and Simpson’s evenness index (**Supp Fig. 1C**). Although significant differences in microbiota richness between male MJD and male WT mice were found at 5 weeks (p = 0.0029), no significant differences at 7 or 13 weeks of age were observed, nor for microbiota diversity or evenness at any time point. Female MJD and female WT mice did not demonstrate any significant differences in all three microbiota diversity indices at 5, 7 or 13 weeks-of-age. (**Supp Fig 1A-C).** This observation aligns with our previous observations where we report no significant difference in microbiome alpha diversity between MJD and WT animals [2].

### MJD development was linked with changes in gut microbiota composition

The gut microbiota of both male and female MJD and WT mice was predominantly composed of the *Bacteroidota*, *Firmicutes* and *Verrucomicrobiota* bacterial phyla, among others. Following analysis of the ratio between the *Firmicutes* and *Bacteroidota* relative abundance, no statistically significant differences were detected between male and female MJD and WT mice at 5, 7 or 13 weeks-of-age (**Supp Fig. 1D-E**).

At the bacterial family level, the gut microbiome of most male and female MJD and WT mice was comprised of the *Muribaculaceae*, *Erysipelotrichaceae*, *Akkermansiaceae*, *Lachnospiraceae* and *Lactobacillaceae* families, (**Supp Fig. 1E).** Upon examining the effect of genotype, specific bacterial families were identified to have statistically different abundances between MJD and WT groups (**Table S1**). For example, the abundance of the *Akkermansiaceae* family was significantly higher in male MJD mice at 5 weeks (p = 0.035) and 7 weeks-of-age (p = 0.031) compared to male WT controls.

The microbiome composition was then analysed at the ASV level, and this revealed significantly different ASVs in the gut microbiota of MJD mice at 5 weeks (**Fig. 3A-B**), 7 weeks (**Supp Fig. 2**) and 13 weeks-of-age (**Supp Fig. 3**) compared to WT control mice. Numerous ASVs in bacterial families, including members of the *Lachnospiraceae, Muribaculaceae* and *Oscillospiraceae* families, displayed divergent abundances in male MJD and female mice compared to male and female WT mice at the three ages examined. Notably, many ASVs in the *Lachnospiraceae* family were seen at a lower abundance in male MJD and female MJD mice, particularly at 7 and 13 weeks-of-age. Many ASVs belonging to the *Muribaculaceae* and *Oscillospiraceae* families also had a lower relative abundance in MJD mice, however, different sets of ASVs in the same families were seen at a higher abundance in MJD mice. Despite the similarities at the family level taxonomic assignment, the exact set of ASVs that were significantly different between MJD and WT mice were age-specific, with no single ASV consistently found to be differentially abundant at all three ages. Further, most differentially abundant ASVs were specific to each sex, where only 3 ASVs, *Muribaculaceae* (ASVs 16 and 18) and *Lachnospiraceae* (ASV 595), showed similar MJD-associated shifts in the microbiome of male and female mice at 5 weeks.

**Fig. 3.**
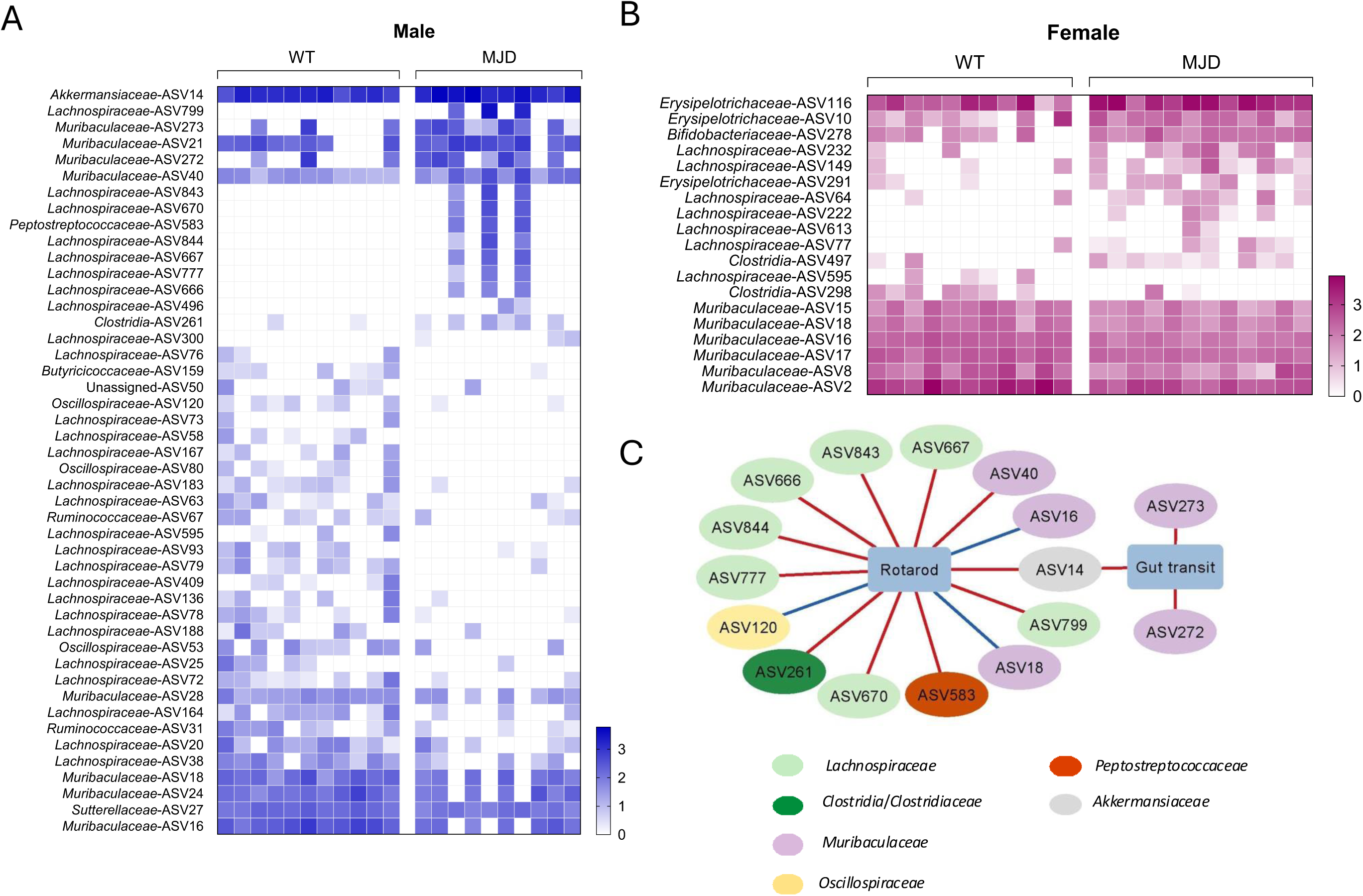
Gut microbiota is different between MJD and WT mice at 5 weeks-of-age, and these shifts correlate with the onset of motor and colonic impairment. Amplicon sequence variants (ASVs) that significantly differed in abundance (p < 0.05) between MJD and WT groups are presented for 5-week-old **(A)** male and **(B)** female mice. Unpaired heatmaps display the relative abundance (Log10 transformed) of each ASV, where rows denote ASVs, and columns denote each mouse. For male mice, blue corresponds to the highest relative abundance, and white corresponds to the lowest abundance. For female mice, pink corresponds to the highest abundance and white corresponds to the lowest abundance. **(C)** Network demonstrating correlations between ASVs that were different between male MJD and WT mice at 5 weeks and their gut transit and rotarod performance at 9 and 11 weeks-of-age, respectively. Pairwise correlation analyses were used to identify associations of statistical significance (p < 0.05) and were used to create networks.

We assessed how changes in the gut microbiome observed in pre-symptomatic (5 weeks) MJD mice correlated with the onset of colonic and motor dysfunction at 9 and 11 weeks-of-age, respectively (**Fig. 3C**). Due to the delayed onset of motor dysfunction and absence of gut function differences in female MJD mice, correlation analyses were performed only on male animals. The relative abundance of two ASVs in the *Muribaculaceae* family positively correlated with gut transit time at 9 weeks, indicating that male mice with a higher abundance of these ASVs in the pre-symptomatic stage later demonstrated a longer gut transit time at 9 weeks-of-age. A single ASV in the *Akkermansiaceae* family showed positive correlations with both gut transit and rotarod performance. This suggest that male animals having a higher abundance of this ASV at the pre-symptomatic stage, later had a longer gut transit time and better rotarod performance. The relative abundance of seven ASVs in the *Lachnospiraceae* family at 5 weeks positively correlated with rotarod measurements at 11 weeks, indicating that pre-symptomatic male mice with a higher abundance of the specific ASVs had better rotarod performance at week 11.

### Protein aggregates are present in the brain of male MJD from 5 weeks-of-age

To gain an understanding of the time course of neurodegeneration within the brain compared to the gut microbiome changes described above, we performed immunohistochemistry on brain tissue obtained from Cohort 2 mice (all male) euthanised at 5, 7, 9, 11 weeks-of -age, and male Cohort 1 mice euthanised at 13 weeks-of-age (**Fig 4A**). Staining for ataxin-3 in sections from the medulla oblongata revealed ataxin-3 aggregates in samples from MJD mice but not WT mice (**Fig 4B**). MJD mice exhibited significantly more ataxin-3 aggregates than WT, indicated by a genotype effect (p < 0.0001), together with an effect of age (p < 0.0001), and an interaction effect (p < 0.0001). Tukey post-hoc analysis revealed that MJD mice had significantly more ataxin-3 protein aggregates that WT mice from 7-weeks-of-age old onwards (p < 0.0001; **Fig 4C**).

**Fig. 4.**
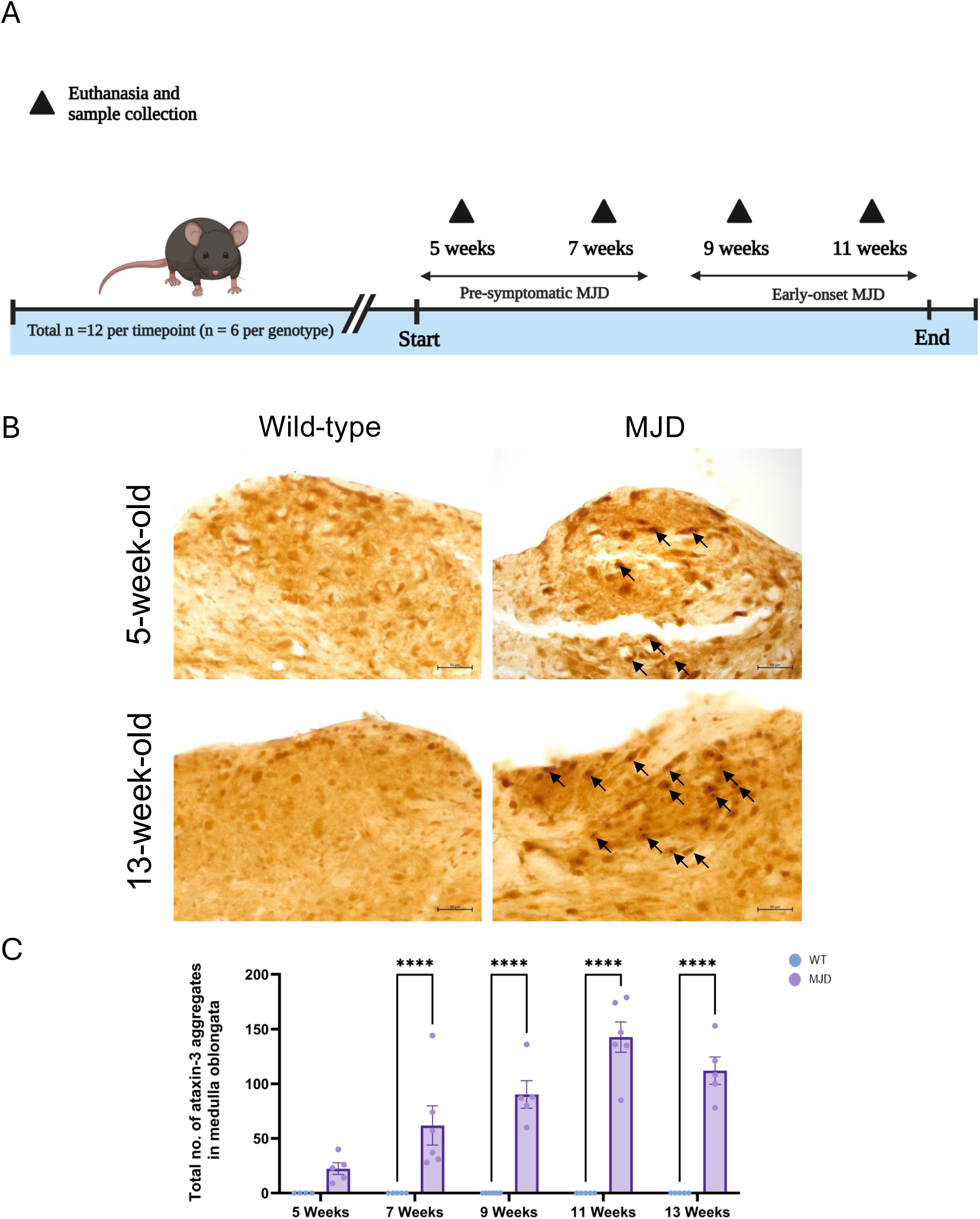
MJD mice develop ataxin-3 aggregates during pre-symptomatic disease stage at 5-weeks-old. **(A)** Experimental timeline of Cohort 2 mice. A total of 48 WT and MJD mice (n = 12 per time point, n = 6 per genotype within each timepoint) were euthanised for sample collection at the indicated timepoints. Timeline figure was created using BioRender.com. **(B)** Immunohistochemical staining of ataxin-3 in sections from the medulla oblongata of pre-symptomatic 5-week-old and early-onset stage 13-week-old WT and MJD mice. Scale bars represent 50μm **(C)** MJD mice exhibited significantly more ataxin-3 aggregates within the medulla oblongata than WT mice across all timepoints. Statistical analysis performed using unpaired t-test for western blotting and two-way ANOVA followed by Tukey’s post-hoc analysis for immunohistochemical staining. All vales represent mean ± SEM. *** p < 0.001, **** p < 0.0001.

### Gut morphology is not altered in male MJD mice

Next, we investigated whether the altered gastric motility was due to altered morphology or neuronal loss within gastrointestinal tissue. To confirm whether h*ATXN3* and human ataxin-3 are present in the gut of MJD mice, we performed RT-PCR and western blotting (respectively) on samples from the small intestine and colon of MJD and WT mice. We observed h*ATXN3* mRNA expression in the colon and small intestine of the MJD mice, but not their non-transgenic littermates (**Fig 5A**). We also detected human mutant ataxin-3 at near-endogenous levels in the small intestine of the mutant animals (**Fig 5B-C**). However, we were unable to find the presence of full-length human mutant ataxin-3 protein in the colon of CMVMJD135 mice (data not shown).

**Fig. 5.**
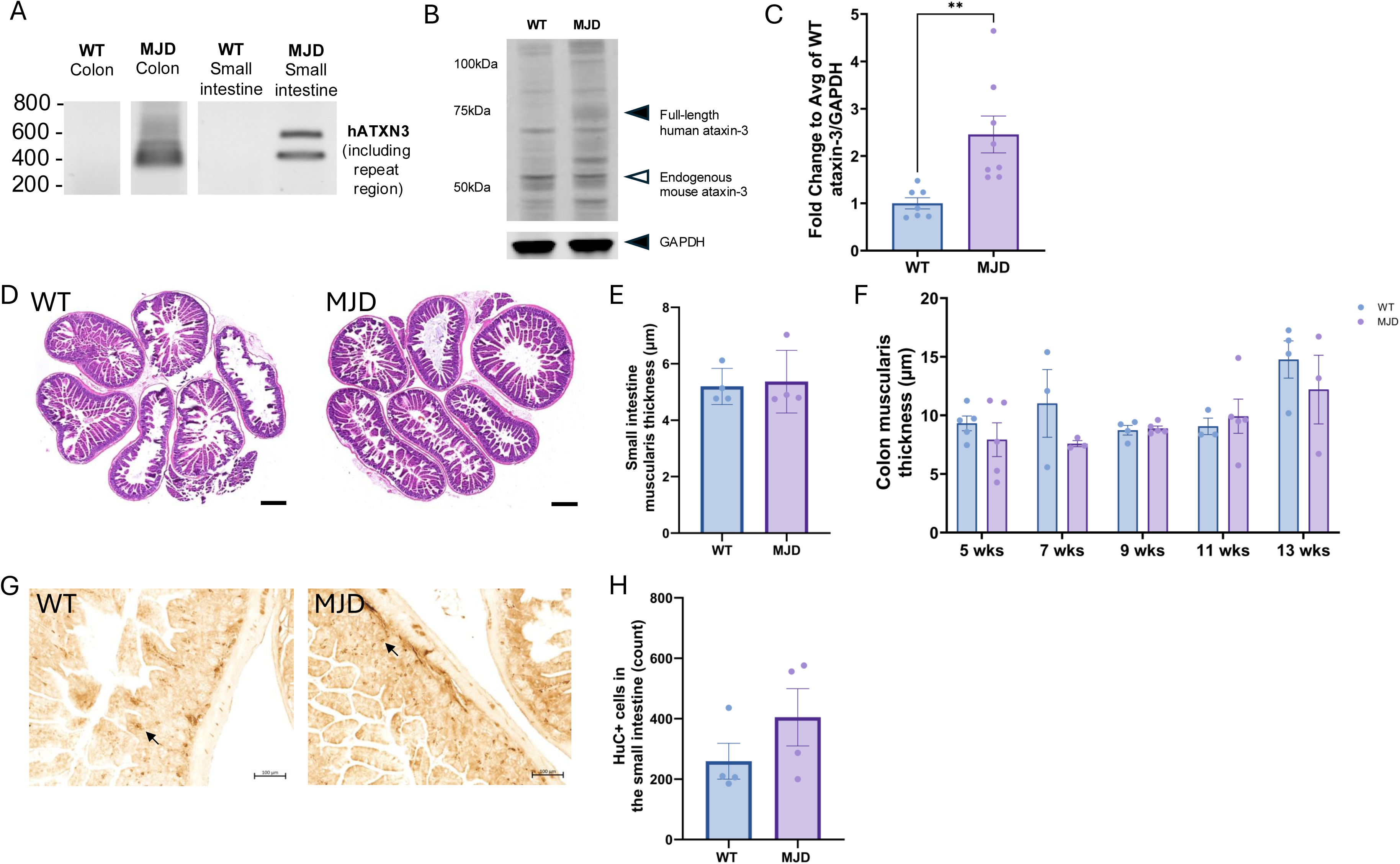
MJD mice do not exhibit altered gut morphology or enteric neuron number, compared to WT mice. **(A)** Representative RT-PCR of non-transgenic wild type (WT) mice and CMVMJD135 (MJD) mice confirms increased expression of human *ATXN3* in the colon and small intestine of 13-week-old MJD mice compared to WT mice. **(B)** A representative blot of WT and MJD mouse small intestine confirms increased expression of full-length ataxin-3 in the small intestine of 13-week-old MJD mice compared to WT mice. **(C)** Quantification of full-length ataxin-3 expression demonstrates a significant increase in (p= 0.0052) in the small intestine of 13-week-old MJD mice compared to WT mice. **(D)** Representative images of H&E staining of WT and MJD small intestine tissue. Scale bars represent 500μm. **(E)** Small intestine muscularis layer thickness quantification of 13-week-old mice revealed no difference in thickness between WT and MJD mice. **(F)** Quantification of colon muscularis layer thickness in the colons of 5, 7, 9, 11 and 13-week-old mice presented no significant effect of age or genotype on thickness. **(G)** Representative images of HuC+ immunolabelling in male 13-week-old WT and MJD mouse small intestine. Scale bars represent 100μm. **(H)** Quantification of HuC+ immunolabelling within small intestine samples of male 13-week-old mice showed no significant differences in total enteric neuron populations between WT and MJD mice. Statistical analysis performed using unpaired t-test for western blotting, small intestine muscularis and HuC analysis. Colon muscularis analysis performed using two-way ANOVA followed by Tukey’s post-hoc analysis. All vales represent mean ± SEM. **p < 0.01

Histochemical staining (haematoxylin and eosin) of formalin fixed paraffin embedded small intestine tissue revealed no detectable differences in the morphology of small intestine tissue. Image analysis revealed no significant difference (p = 0.7997) in the thickness of the muscularis layer of the small intestine of male WT and MJD mice (at 13-weeks-old) (**Fig 5D-E**). Similarly, no significant effect of genotype (p = 0.1661) was found in the thickness of the colon muscularis of male MJD and WT mice at 5, 7, 9, 11 or 13 weeks old (**Fig 5F)**, however, we found a significant effect of age (p = 0.0123).

To confirm whether the altered gastric motility we identified was due to altered neural innervation within the gastrointestinal tract, we performed immunostaining in small intestine tissue of 13-week-old male mice for HuC as a marker of enteric neurons (**Fig 5G**). We found no significant difference (p = 0.2409) in total enteric neuron numbers per animal (**Fig 5H**).

### Altered abundance of gastric endocrine factors and inflammatory markers present in the small intestine of male MJD mice

A major modulator of gut motility is the release of endocrine factors such as cholecystokinin (*Cck*) and ghrelin (*Ghrl*), among others. We performed qPCRs to compare the expression of these genes within the small intestine tissue of our 13-week-old male WT and MJD mice and found that male MJD mice had increased *Cck* (p < 0.01, **Fig 6A**), and *Ghrl* mRNA abundance (p < 0.01, **Fig 6B**). We also performed RT-qPCRs to compare the abundance of inflammatory modulators in the small intestine tissue and observed increased abundance of heme oxygenase (*Ho1*) (**Fig 6D**) and decreased inducible nitric oxide synthase (*Nos2*) mRNA (**Fig 6F**) in male MJD mice. Whilst no significant differences were found in the abundance of tumour necrosis factor alpha (*Tnfa*) mRNA (p = 0.6532, Fig 6G), Complement C1q A Chain (*C1qa*) mRNA (p = 0.2579, **Fig 6C**), an increased abundance of interleukin-1 beta (*Il1b*) mRNA was found in the small intestines of male MJD mice, compared to WT controls (p < 0.01, **Fig 6G**).

**Fig. 6.**
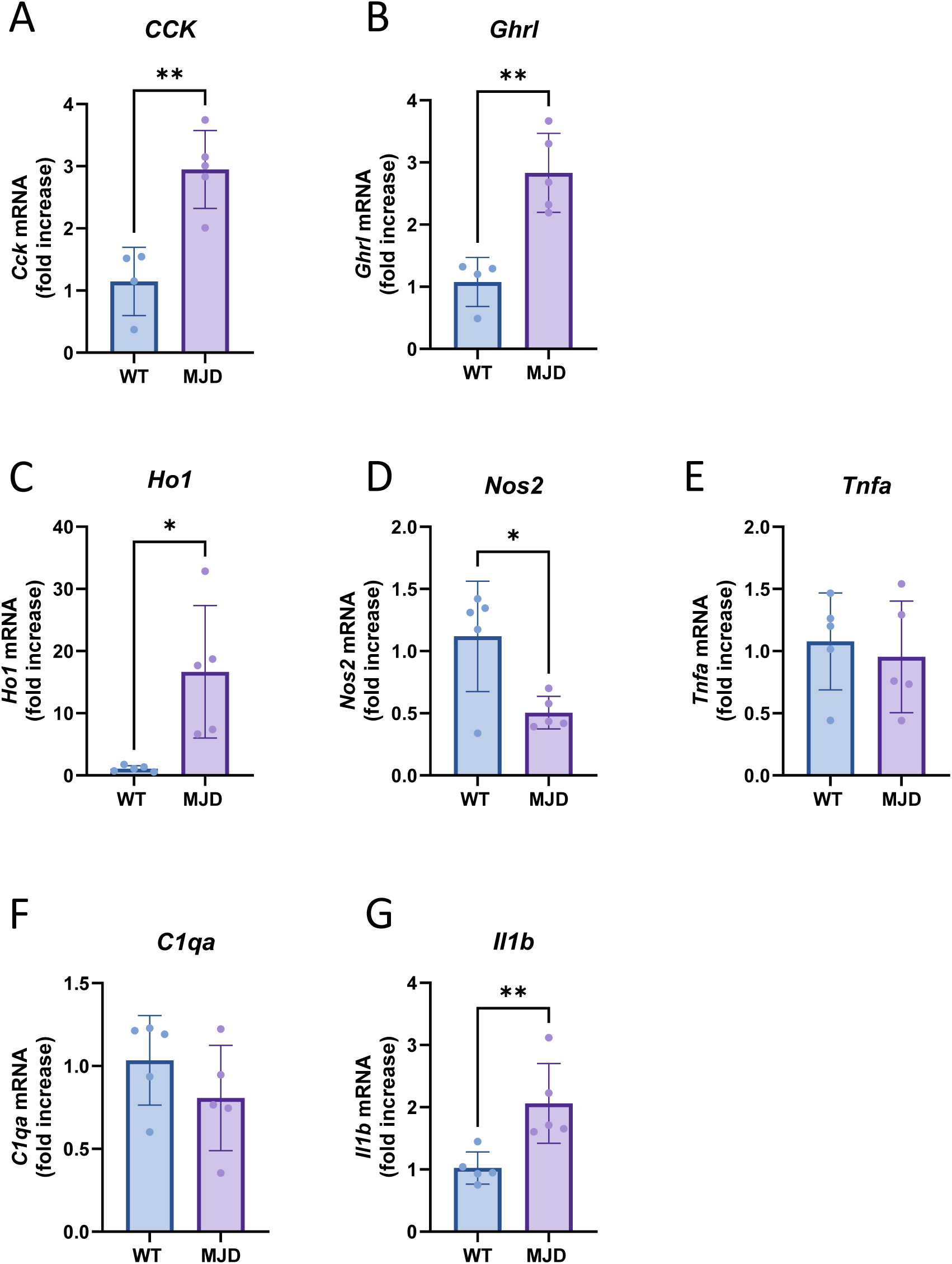
Altered expression of genes involved in the regulation of gastrointestinal function and inflammation within small intestine samples of 13-week-old MJD versus WT mice. Gene expression analysis via RT-qPCR showed that (A) *Cck* and (B) *Ghlr* mRNA levels were significantly increased in MJD mice compared to WT mice. MJD (n = 5) and WT (n = 4). No significant differences were found in the mRNA expression levels of inflammation mediators *C1qa* (C) *Il1b* and (D) *Tnfa* whilst a significant difference was observed in the expression levels of (E) *Ho1* and (F) *Nos2*. The reference gene used in these analyses was Ribosomal Protein L-27 (*Rpl27*). MJD (n = 5) and WT (**A-B**: n = 4 and **C-F**: n = 5). Statistical analysis was performed using an unpaired t-test. All values represent mean ± SEM. *p < 0.05, **p < 0.01.

### No difference in faecal butyrate or abundance of malabsorbed fat in colonic faecal samples of 11-week-old mice

To examine whether the fast gut transit phenotype was related to altered abundance of the short-chain fatty acid butyrate in the MJD mice, we performed gas chromatography mass spectroscopy (GCMS) analysis to quantify the butyrate levels within faecal samples collected from the mice at 11 weeks-of-age. This analysis found similar levels of butyrate in the faecal samples of male MJD and WT mice (**Supp. Fig. 4A**).

To examine whether the fast gut transit phenotype that we identified was affecting functions such as fat absorption, we examined faecal samples from the 11-week-old male WT (n = 6) and MJD (n = 6) mice from Cohort 2. We performed Sudan III staining for malabsorbed faecal fat on the samples. Quantification of malabsorbed faecal fat showed no difference in the total amount present between the WT and MJD mice (**Supp. Fig. 4B-C**).

## DISCUSSION

### Colonic dysfunction occurs early in disease of male MJD mice

Our study demonstrates for the first time that male transgenic MJD mice develop altered colonic function, particularly in gut transit time. A reduction in the time taken for food to traverse the gastrointestinal tract was observed in male MJD mice as early as 9-weeks-of-age, which became more profound as the disease progressed. Interestingly, the onset of this faster gut transit in male MJD mice was identified before detecting impaired performance on the accelerating rotarod, evident by 11 weeks-of-age. While significant differences in faecal output measures between male MJD and male WT mice were also observed at 13 weeks-of-age, it is difficult to propose the age of onset as tests were only conducted once.

Our study also confirms that MJD is associated with changes in the structure and composition of the gut microbiome, where these shifts initiated at pre-symptomatic stages in male MJD mice as early as 5 weeks-of-age. To date, the earliest gut microbiome changes reported in MJD mice were at 7-weeks-of-age, which occurred prior to the onset of any known neurological and motor impairment of the disease [2]. We report here that MJD-associated gut microbiome changes occur even earlier, by 5-weeks-of-age in male MJD mice, and that the gut microbiome becomes increasingly divergent by 13-weeks-of-age. These gut microbiota changes occur before the onset of altered gut transit times at 9 weeks and motor impairment at 11-weeks-of-age. Our study is the first to report that functions of the gut and gut microbiota change with the development of MJD and that gut microbiome changes occur prior to the onset of functional colonic changes and motor impairment. This suggests that colonic dysfunction, as well as even earlier changes in the gut microbiome, are among the first changes to emerge and potentially contribute to the development of MJD related characteristics.

Gut microbiota changes and colonic dysfunction have also been shown to occur concurrently in other neurodegenerative diseases such as PD [20, 38, 39], with microbial shifts hypothesised to trigger changes in colonic function and vice versa. However, constipation, decreased intestinal motility (i.e., slower gut transit) and decreased faecal output are frequently reported in examined neurodegenerative disease [1, 40–43]. Our study demonstrates a paradigm of the opposite effect, showcasing faster gut transit and increased faecal output in our male MJD mice, with no significant changes in female MJD mice.

Alterations in the gut microbiome have been previously reported in patients and in mouse models of MJD [2], PD [12], Alzheimer’s disease (AD) [7] and amyotrophic lateral sclerosis (ALS) [44] compared to controls. Changes in the abundance of specific gut microbes have also been linked with the extent of disease severity in AD [45, 46] and ALS [47]. Our previous work investigating gut dysbiosis in CMVMJD135 mice modelling MJD reported that disease characteristics were exacerbated when the gut microbiota during the pre-symptomatic stage maintained a lower abundance of specific ASVs in the *Lachnospiraceae*, *Oscillospiraceae*, *Rikenellaceae* and *Ruminococcaceae* families [2]. The study reported that specific ASVs from these families in pre-symptomatic male and female mice demonstrated positive correlations with rotarod duration and negatively correlated with neurological score during well-established disease stages [2]. Other studies using AD mouse models have reported *Muribaculaceae* to contribute to the pathophysiology of postoperative neurological dysfunction, as well as a decreased abundance of this family in ileitis and T-cell-induced colitis [48–50]. Due to this, these associations require biological validation using targeted assays, for instance by administering bacterial isolates and examining the impact on the development and pathology of MJD.

### Male MJD mice with colonic dysfunction showed no changes in intestinal morphology

It can be speculated that colonic dysfunction in MJD mice may stem from dysregulated autonomic control of the gastrointestinal tract due to neurodegeneration of autonomic nuclei. However, as our study identified faster gut transit and increased faecal output in our male MJD mice, perhaps an effect of neurodegeneration on the enteric and autonomic neurons may not be at play. Indeed, we found that the number of enteric neurons present within the small intestine of male MJD mice did not significantly differ to the number detected in the small intestine of male WT controls. Furthermore, no signs of ataxin-3 aggregates were found in the small intestine or colon of MJD or WT controls. We also examined the thickness of the muscularis layer of the small intestine and colon of 9-week-old male WT and MJD mice, with no differences found. These findings suggest that changes in colonic motility and faecal output are not a result of morphological changes in these tissues.

### Gut microbiome may potentially drive colonic dysfunction in male MJD mice

We report gut microbiome changes in MJD mice that precede colonic dysfunction and motor impairment. Gut microbiome changes were present at pre-symptomatic stages in 5-week-old male MJD mice and were correlated with the severity of gut dysfunction and motor impairment present in the mice at symptom onset stages. These findings indicate that some gut microbiota changes are predictive of the extent of functional colonic and motor impairment in MJD. For example, the abundance of ASVs in the *Lachnospiraceae* and *Muribaculaceae* families during pre-symptomatic stages positively correlated with rotarod and gut transit at 9 and 11-weeks-of-age, respectively, indicating a link between these ASVs with improved motor and colonic function.

We previously reported that disease characteristics were exacerbated when the pre-symptomatic gut microbiota carried a lower abundance of specific ASVs in the *Lachnospiraceae*, *Oscillospiraceae*, *Rikenellaceae* and *Ruminococcaceae* families in male CMVMJD135 mice. The study reported that specific ASVs from these families in pre-symptomatic male and female mice demonstrated positive correlations with rotarod duration and negatively correlated with neurological score during well-established disease stages [2]. Other studies have reported *Muribaculaceae* to contribute to postoperative neurological dysfunction in mice, and decreased abundance of this family has been reported in ileitis and T-cell-induced colitis [48–50]. Certainly, it has been known for over fifty years that germ-free mice develop distended ceca, slowed intestinal transit and gastric emptying [51, 52]. Husebye *et al.* have demonstrated that this delay could be partially reversed by colonization with *Lactobacillus acidophilus* and *Bifidobacterium bifidum* [53]. Nevertheless, the potential microbial role in altered gut transit, reported here, requires biological validation using targeted assays, for instance, by administering bacterial isolates and examining the impact on measurable readouts.

### Potential mechanisms of fast gut transit in male MJD mice

It is plausible that changes to the gut microbiome at early ages of disease may drive changes in metabolites or immune signalling, producing an increased transit time as reported in the present study. Gastrointestinal motility is known to be modified by changes in the levels of inflammatory mediators, such as immune cells and cytokines [54], in diseases such as irritable bowel syndrome [55]. SCFAs, produced by the gut microbiome, have been reported to influence microglial function (e.g., maturation and activation) within the CNS and may, therefore, indirectly contribute to neuroinflammation and neurodegeneration [56]. In our study, we did not identify any alteration to faecal butyrate levels, or signs of fat malabsorption, suggesting that those pathways are unlikely to be involved in the reported changes.

RT-qPCR analysis of samples from the small intestine of the mice did identify increased abundance of the pro-inflammatory cytokine *Il1b* mRNA in male MJD mice compared to WT controls. Immune cells within the intestinal mucosa are known to be producers of IL-1β, triggered by stimuli such as injury and dysbiosis [57]. In fact, IBD patients have elevated levels of IL-1β in their intestinal tissue, with increased intestinal inflammation correlated with increasing levels of intestinal IL-1β [58].

We also identified increased abundance of *Ho1* mRNA and decreased *Nos2* mRNA in the small intestines of the male MJD mice. HO-1 is an enzyme released in response to cell stressors such as oxidative stress and inflammation, acting as an anti-inflammatory molecule, such as intestinal injury, inflammatory bowel disease, or exposure to toxins [59]. Studies on vascular smooth muscle cells have reported that IL-1β can lead to increased *HO1* mRNA after 24 hours of administration [60]. Furthermore, they identified that this increase was due to an elevation in the rate of *HO1* gene transcription. On the other hand, we observed a decrease in the mRNA levels of iNOS producer *Nos2* in MJD mice compared to WT. Although it is well-documented that iNOS increases during inflammation [61], several studies have demonstrated the possible existence of a negative feedback mechanism where increased nitric oxide (NO) during inflammation leads to decreased *Nos2* mRNA levels, suggesting a regulatory loop that prevents excessive NO production and limits further inflammation [62–64]. Furthermore, up-regulation of *HO1* in astrocytes is associated with down-regulation of *NOS2* expression and thereby NO production [60], and elevated HO-1 and decreased NO have each been reported to produce fast gut transit [65, 66].

Furthermore, we examined changes in gut hormones, CCK and ghrelin, which are peptides known to regulate gastrointestinal motility, appetite, and digestion [67]. These hormones were selected due to their critical roles in regulating gut function, which we observed were disrupted in disease. Specifically, CCK has been implicated in slowing gastric emptying and affecting motility in the small intestine, while ghrelin is known to accelerate gastric motility and promote food intake [68]. Altered levels of these hormones could provide insight into the underlying mechanisms of gastrointestinal dysfunction in MJD.

Our study revealed that male MJD mice at 13 weeks-of-age exhibited significantly higher levels of *Cck mRNA* and *Ghrl mRNA* in the small intestine. This time point coincides with the onset of faster faecal transit in these mice, suggesting a potential role for these hormones in the observed gastrointestinal abnormalities. Given that the mutant mice exhibited a faster food transit, we expected a decrease in CCK levels in the disease model. However, it is important to note that our study measured mRNA levels of these hormones and thus, it remains unknown whether hormone levels in the blood are altered. Additionally, the relation between CCK levels and gastric emptying seems to depend on other physiological conditions. For example, Bento da Silva *et al* observed that slower gastric emptying induced by physical exercise in rats was associated with a decrease in circulating CCK levels [69]. However, they also found that this decrease in blood CCK correlated with an increase in *Cck* mRNA expression in the pylorus, suggesting that CCK regulation at the transcriptional and protein levels may be differentially controlled. This highlights the complexity of CCK’s role in gastrointestinal physiology, where its local expression and systemic effects may be influenced by various physiological conditions.

Remarkably, increased ghrelin levels have been shown to correlate with faster gastric emptying in pathophysiological conditions, such as diabetes and obesity, where gut motility is altered [70]. Moreover, ghrelin has been recognised for its protective properties against stressors, such as 1-methyl-4-phenyl-pyridinium (MPP+) [71, 72]. Studies have shown that ghrelin can mitigate damage from oxidative stress and inflammation, possibly through activation of the Nrf2 pathway and the subsequent upregulation of *HO1* [68].

Interestingly, both CCK and ghrelin levels have been associated with gut microbiota activity. For example, exposure to butyrate stimulates ghrelin secretion from 2D human enteroids [73]. Mice with fructose malabsorption have increased density of I-cells and *Cck* mRNA expression in the ileum and cecum, which is also accompanied by increased levels of *Actinobacteria*, *Bacteroidetes*, and *Lactobacillus johnsonii* [74]. Furthermore, levels of cecal propionate are increased in mice with fructose malabsorption and propionate can trigger *Cck* gene expression *in vitro*, suggesting it may modulate CCK levels in mice [75]. These findings suggest a strong relationship between gut microbiota composition, gut hormone regulation, and gastrointestinal function. The interplay between microbial metabolites, such as short-chain fatty acids, and hormone secretion highlights the potential role of the gut microbiome in modulating CCK and ghrelin levels, which in turn may influence gut motility, inflammation and disease progression. Further studies, including protein and hormone release assays, would be necessary to confirm whether the observed mRNA changes correspond to altered peptide release and its functional consequences.

### MJD mice exhibited sex-specific differences in colonic dysfunction, gut microbiome and motor symptom onset

We have previously reported sex-dependant development of motor symptoms and gut microbiome alterations in male and female MJD mice. Interestingly, we have found that in a similar manner to the sex difference in the development of motor symptoms in CMVMJD135 mouse, male MJD mice also develop gut microbiome and gut function changes prior to female MJD mice. We report sex differences not only in gut microbiome alterations and onset of motor symptoms but a lack of colonic dysfunction in female MJD mice until 13-weeks-of-age, which in contrast was evident in male MJD mice as early as 9-weeks-of-age. Similar findings have been reported in a rodent model of Huntington’s disease, with Kong *et al*. reporting a sex difference in the gut microbiota and faecal output of Huntington’s disease mice [76]. However, in their study they only profiled the faecal microbiota at a singular age and only during a pre-symptomatic disease stage [76]. Our study provides further insights into sex differences in CMVMJD135 mice modelling MJD and neurodegeneration more broadly.

Sex-driven variations linked with neurodegenerative disease are potentially associated with sex hormones and their neuroprotective effects or mechanisms [77, 78]. It can be postulated that sex hormones may affect the pathophysiology of MJD through the compositional modulation of the gut microbiome and/or inflammatory and hormonal changes, transmitted from the colon to the central nervous system (CNS) through the brain-gut-microbiome (BGM) axis [79, 80]. Future work is required to confirm the associations between sex hormones, gut microbiota changes and the development of colonic dysfunction and motor impairment in MJD.

## Conclusions

This is the first report of the occurrence of colonic dysfunction in the development of MJD. It is also the earliest report of gut microbiota changes in the development of MJD. In conjunction to changes in colonic function and the gut microbiota associated with MJD, we report that these changes occur during early-onset and pre-symptomatic disease stages, respectively, and prior to the onset of any motor dysfunction in male MJD mice compared to male WT controls. We find that the presence of ataxin-3 protein aggregates in the brain and altered composition of the gut microbiota are the earliest phenotypes to occur in the male MJD mice, present as early as 5-weeks-old. Therefore, these changes may be the trigger of the observed functional changes, including fast gut transit, and motor impairment. Nevertheless, both the altered gut microbiota and colonic dysfunction changes increased during the disease progression in male MJD mice and therefore warrant investigation as potential targets for MJD treatment.

**Supp Fig 1.**
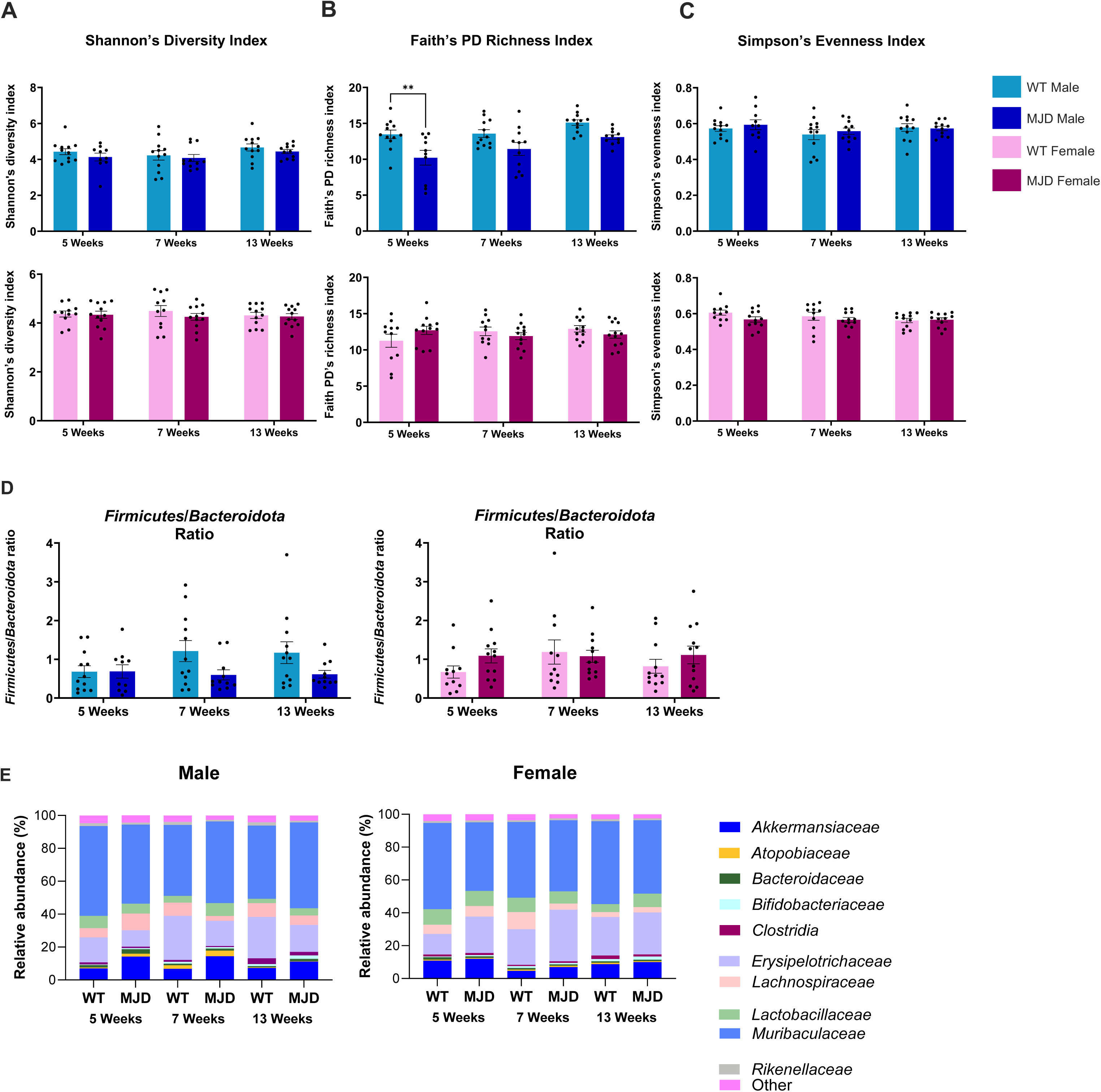
The gut microbiota alpha diversity is shown as **(A)** Shannon’s diversity index, **(B)** Faith’s PD richness index and **(C)** Simpson’s evenness index. **(D)** The ratio between the abundance of the *Firmicutes* and *Bacteroidota* phyla identified in male and female MJD and WT mice at 5, 7 and 13-weeks-of-age. Data is shown as mean ± SEM, with significance determined using a two-way analysis of variance (ANOVA) and Šídák’s multiple comparisons test. **(E)** Family level relative abundance (%) and taxonomic composition of the gut microbiota of faecal samples collected from male and female MJD and WT mice at 5, 7 and 13-weeks-of-age. Families with a relative abundance < 1% in all samples are categorised as ‘Other’.

**Supp Fig 2.** Amplicon sequence variants (ASVs) with significantly different (p < 0.05) abundances between MJD and WT groups are presented for 7-week-old **(A)** male and **(B)** female mice. Unpaired heatmaps display the relative abundance (Log10 transformed) of each ASV, where rows denote ASVs, and columns denote each mouse. For male mice, blue and white correspond to the highest and lowest relative abundance, respectively. For female mice, pink and white corresponds to the highest and lowest abundance, respectively.

**Supp Fig 3.** Amplicon sequence variants (ASVs) that showed significant differences between MJD and WT **(A)** male and **(B)** female at 13-weeks-of-age. The relative abundance (Log10 transformed) for each ASV is shown in the unpaired heatmaps. Rows denote ASVs, and columns denote each mouse. Blue and white in the heatmap for male animals correspond to the highest and lowest relative abundance, respectively, whereas pink and white corresponds to the highest and lowest abundance, respectively in the heatmap for females.

**Supp Fig 4. Altered faecal butyrate and malabsorbed faecal fat are not present in the MJD mice at 11 weeks old. (A)** Gas chromatography mass spectrometry (GCMS) was utilised to detect butyrate concentration within faecal samples collected from the male and female mice (WT and MJD) at 11 weeks old (n = 12 female WT, n = 12 female MJD, n = 12 male WT, n = 11 male MJD), but no difference was found between the experimental groups. **(B)** Quantification of fat malabsorption in colonic faecal samples of 11-week-old mice through use of a Sudan III stain for malabsorbed faecal fat in colonic faecal samples from 11-week-old WT and MJD mice. Scale bars represent 50μm **(C)** Quantification of malabsorbed faecal fat showed no difference in the total amount present between WT (n = 6 mice) and MJD (n = 6 mice) mice (p = 0.9790). Statistical analysis performed using an unpaired t-test. All values represent mean ± SEM.

**Supp Table 1.** The list of bacterial families with significantly different relative abundances between MJD and WT animals. LEfSe analysis between genotypes, per sex at each of the three timepoints were conducted separately. Whether a specific family is higher in MJD or WT is shown with the exact group demonstrating a higher abundance listed under the column Group (M-male, F-female, 5, 7 or 13 denote the age). The LDA score and significance is shown for each of the distinctly abundant bacterial families.

## Supplementary Methods

### Gas chromatography mass spectrometry for detecting butyrate concentration within faecal samples

Butyrate was extracted from faecal samples collected from male mice at 11-weeks-of-age. 40mg of faecal sample from each animal was mixed with acetonitrile and homogenized. The resulting homogenate was centrifuged (3,000 x g, 10 min at 4 °C) and supernatant (100 µL) derivatized together with 10 µL of deuterated –d7-butyric acid and 10 µL of 40 % (v/v) of Pentafluorobenzyl Bromide acid (PFBBr) (cat# TS-58220, Thermo Fisher Scientific) at 60 °C for 1 hr. The derivatized samples were then loaded into the GCMS machine. Acetonitrile was used as a blank sample to correct the background.

GCMS analysis was performed by selective ion monitoring (SIM) mode in a GCMS -TQ 8030 (Shimadzu, USA) with a SH-17Sil MS column (30 m x 0.25 mm x 0.25 µm thickness) (Shimadzu Part Number: 221-75916-30). The ion source and interface temperature were set at 230 °C and 280 °C, respectively. The linear velocity of helium was kept at 60 cm/s. 1 µL of derivatised sample was injected with 4 min of solvent cut-off time, and the mode of inlet was spitless. The initial column oven temperature was set at 35 °C for 3 min and the increased to 130 °C at the rate of 40 °C/min, then finally 295 °C at 20 °C/min and kept at this temperature for 5 min. The ionisation was carried out in the electron impact (EI) mode at 0.2 kV.

The mass spectrometry data was acquired in selective scan mode using target ions having a mass-to-charge ratio (*m/z*) or 268 and 275 for butyric acid and the internal standard, respectively, with an acquisition event time 0.030 s. Data was processed using LabSolutions GCMS solution version 4.53 (Shimadzu, USA). The average of the area response ratio (butyric acid/internal standard) was used to calculate the concentration of butyric acid in the sample, by plotting against a standard curve.

### Faecal fat staining and quantification

Colonic faecal samples (0.05 g) from 11-week-old WT (n = 6) and MJD (n = 6) mice were homogenised in 50 µL of sterile water with a Dounce homogeniser. 60 µL of 35 % glacial acetic acid and 60 µL of 95 % Sudan III dye (w/v, solubilised in 100 % ethanol) were added to the homogenised mixture and briefly vortexed prior to centrifugation at 1000 rpm for 5 min at 4 °C. 35 µL of the supernatant was collected and transferred onto a microscope slide, then cover slipped with a 22 mm x 22 mm coverslip. Cover slipped samples were heated directly over a flame to boiling point three times, and the slide allowed to cool on ice between each boiling step. Coverslips were sealed with clear nail polish and left to dry overnight in a humid staining chamber at 4 °C prior to imaging.

Nine regions measuring 3000 µm x 3000 µm were sampled from each coverslip, with the regions distributed in a manner that allowed representation from all parts of the coverslip. Sampled regions were isolated using the crop function on the OlyVia software (Olympus) and subsequently analysed using ImageJ. Sudan III dye typically stains for all fat within samples however, fats present due to malabsorption stain red, whilst neutral fats stain yellow. Therefore, to quantify fats present due to malabsorption, red lipid droplets were manually quantified using the Multi-Point counting tool on ImageJ. The sum of the red lipid droplets from the nine regions was taken as the total number of malabsorbed fat for each animal. Data analysis was performed on GraphPad Prism (Version 10) using an unpaired t-test.

