## Supplementary figures for "Gut transit and gut microbiome changes occur prior to the onset of motor impairment in a mouse model of Machado-Joseph disease"

| MJD male vs WT male | | | | |
|---|---|---|---|---|
| | **Family** | **Group** | **LDA score** | **p-value** |
| **Higher in MJD** | *Akkermansiaceae* | M5CMV | 4.931 | 0.035 |
| | *Akkermansiaceae* | M7CMV | 4.667 | 0.031 |
| **Higher in WT controls** | *Clostridia* | M7WT | 4.133 | 0.027 |
| | *Rikenellaceae* | M13WT | 3.988 | 0.010 |

| MJD female vs WT female | | | | |
|---|---|---|---|---|
| | **Family** | **Group** | **LDA score** | **p-value** |
| **Higher in MJD** | *Bifidobacteriaceae* | F5CMV | 4.037 | 0.036 |
| | *Erysipelotrichaceae* | F5CMV | 4.764 | 0.036 |
| | *Lactobacillaceae* | F13CMV | 4.537 | 0.013 |

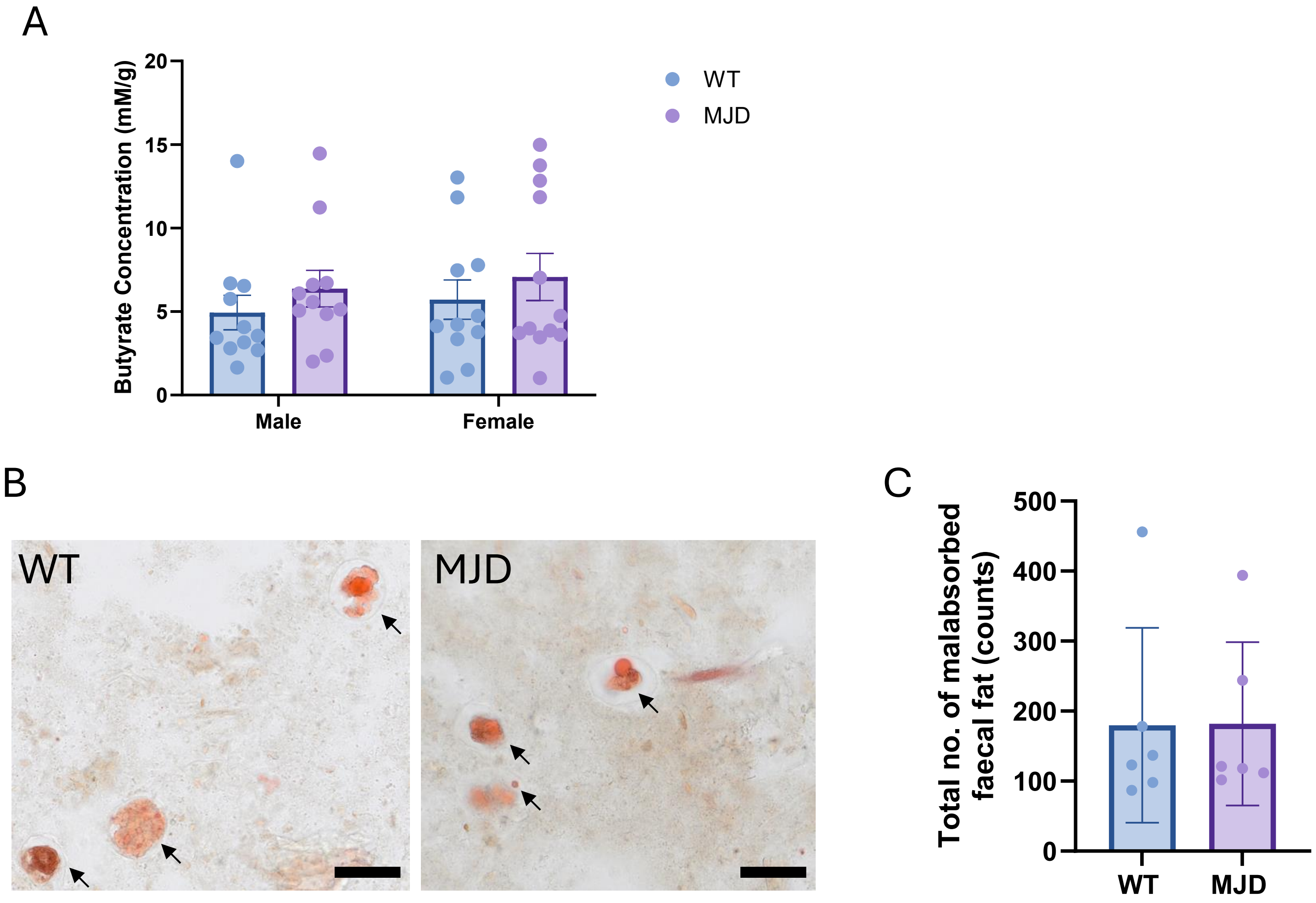
A
Butyrate Concentration (mM/g)
20
15
10
5
0
Male
Female
WT
MJD
B
WT
MJD
C
Total no. of malabsorbed faecal fat (counts)
500
400
300
200
100
0
WT
MJD

### Ataxin-3 COLON AND SMALL INT. PCR WHOLE GEL (Supplementary figure)

| | | | | | | | | | | | | | | | |
|---|---|---|---|---|---|---|---|---|---|---|---|---|---|---|---|
| WT Brain #60 | WT Colon #60 | MJD Brain 6mth old male #1085 | MJD Colon 6mth old male #1085 | WT 13wk old male small intestine #1238 | MJD 13wk old male small intestine #1221 | WT 13wk old male small intestine #1243 | MJD 13wk old male small intestine #1237 | WT 13wk old male small intestine #1281 | MJD 13wk old male small intestine #1250 | WT 13wk old male small intestine #1282 | MJD 13wk old male small intestine #1283 | WT 13wk old male small intestine #1322 | MJD 13wk old male small intestine #1312 | Positive control hATXN3 84Q Plas mid | Negative control MilliQ water |

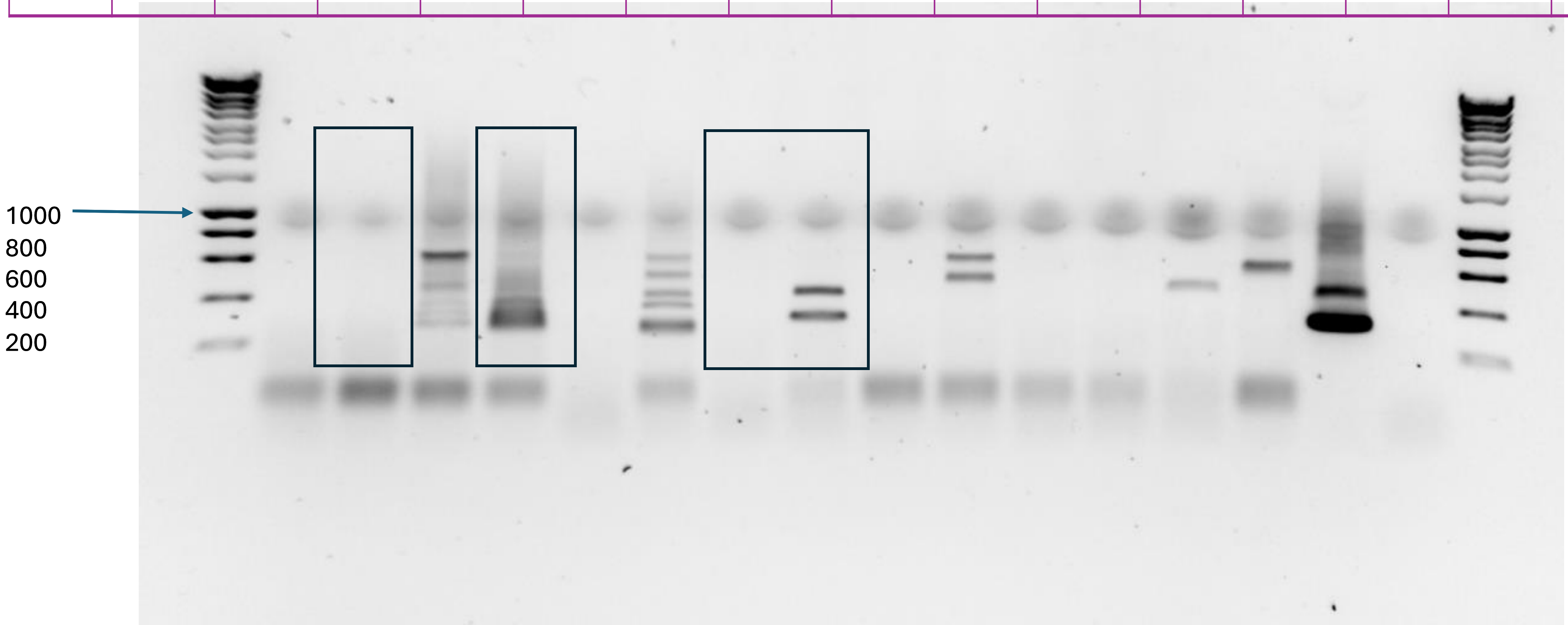

**Ataxin-3 & PolyQ SMALL INT WESTERN BLOT WHOLE GEL – part 1 (Supplementary figure)**

### 20240514 13wk VZ Small Intestine w/ more sample ATXN3 (800nm) & polyQ (680nm)

Potential ATXN3 smear on previous ATXN3 PVDF blot: 1283, 1312, 1321, 1324(very dark)
More samples (newly homogenised) *italicised*

| *WT1* | *MJD1* | *WT1* | *MJD1* | WT1 | *MJD1* | WT1 | MJD1 | SB | SB | *WT1* | MJD1 | WT1 | MJD1 | WT1 | MJD1 | MJD1 |
|---|---|---|---|---|---|---|---|---|---|---|---|---|---|---|---|---|
| *238* | *221* | *243* | *237* | 281 | *250* | 282 | 283 | | | *313* | 312 | 314 | 321 | 322 | 324 | 325 |
| *13wk* | *13wk* | *13w* | *13w* | 13w | *13w* | 13w | 13w | | | *13w* | 13w | 13w | 13w | 13w | 13w | 13w |
| *SI* | *SI* | *k SI* | *k SI* | k SI | *k SI* | k SI | k SI | | | *k SI* | k SI | k SI | k SI | k SI | k SI | k SI |

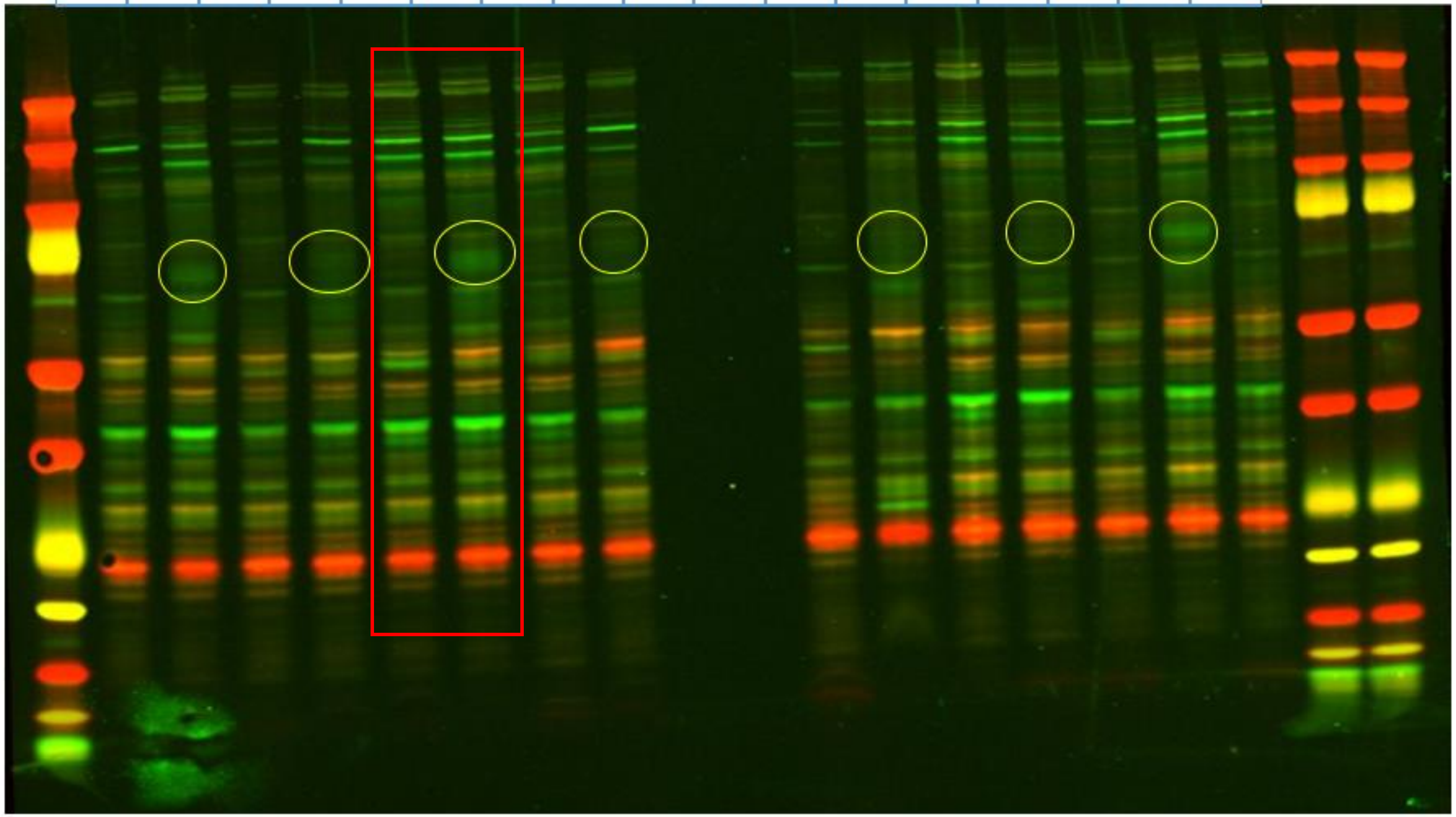

Paulson ATXN3 1:20k;
PolyQ Millipore
1:1000

**Ataxin-3 & PolyQ SMALL INT WESTERN BLOT**
**WHOLE GEL – part 2**
**GAPDH (Supplementary figure)**

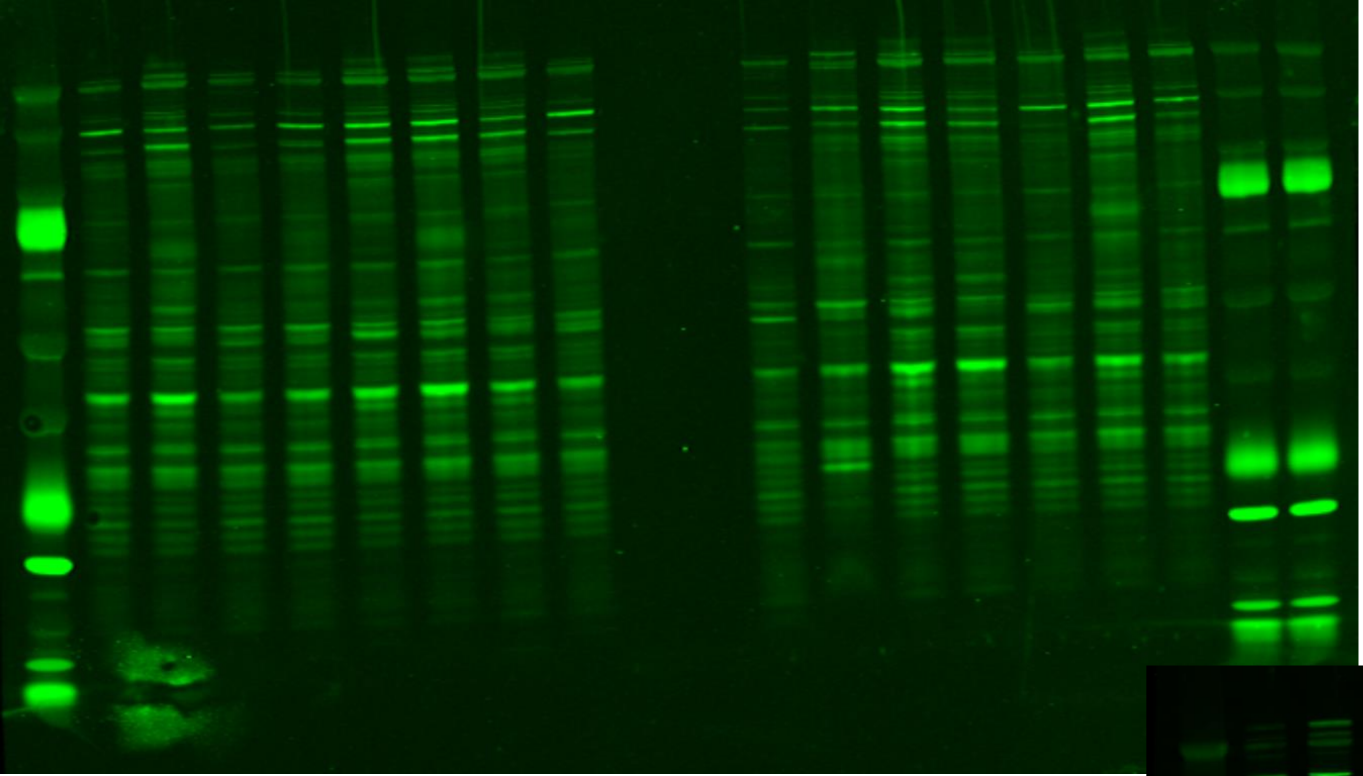

Rb800 ATXN3 (pre-GAPDH)

Rb800 ATXN3 & GAPDH (post GAPDH) - brightness & contrast exaggerated to show both ATXN3 & GAPDH

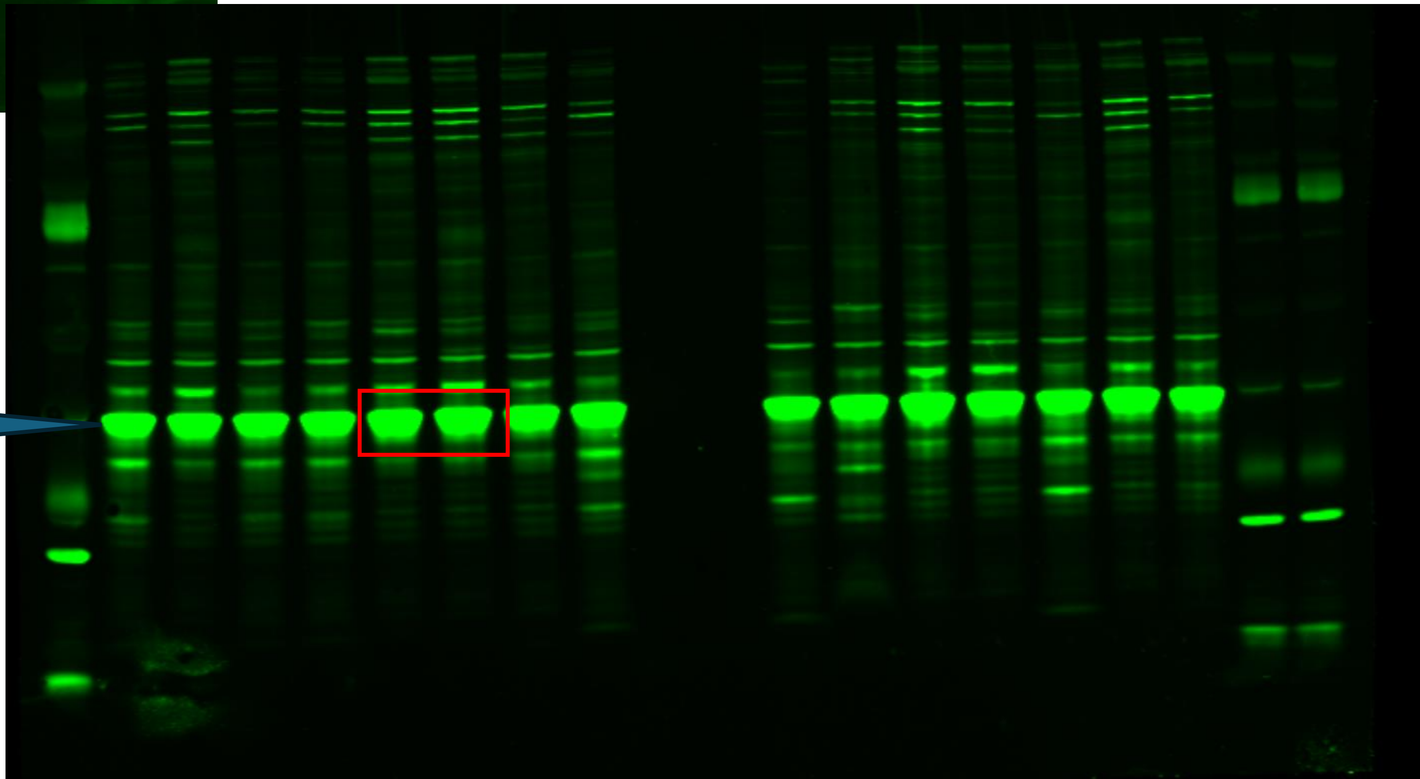

**Ataxin-3 & PolyQ SMALL INT WESTERN BLOT WHOLE GEL – part 3 (Supplementary figure) B&W of Pt 1**

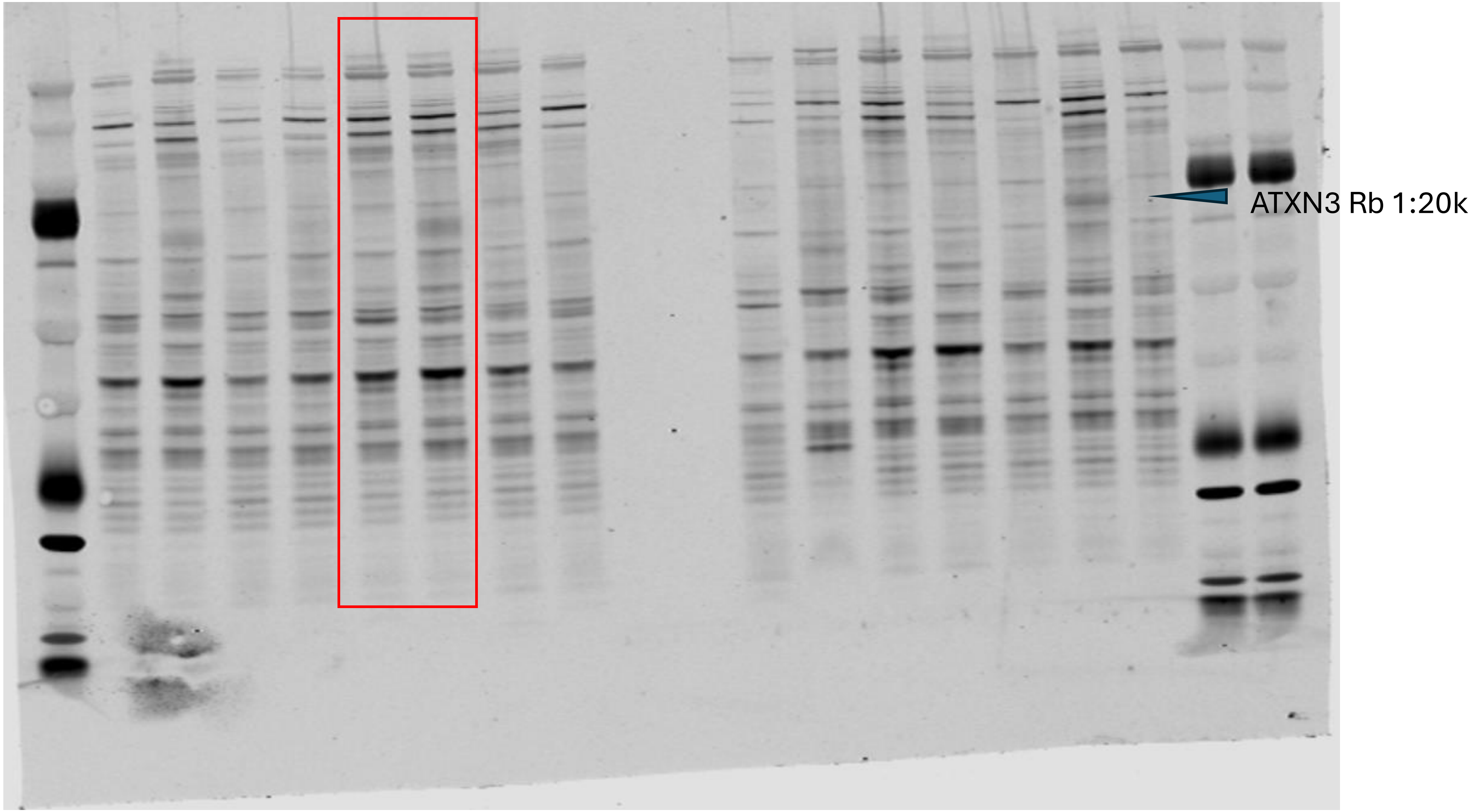

**Ataxin-3 & PolyQ SMALL INT WESTERN BLOT WHOLE GEL – part 4 (Supplementary figure) B&W of Pt 2**

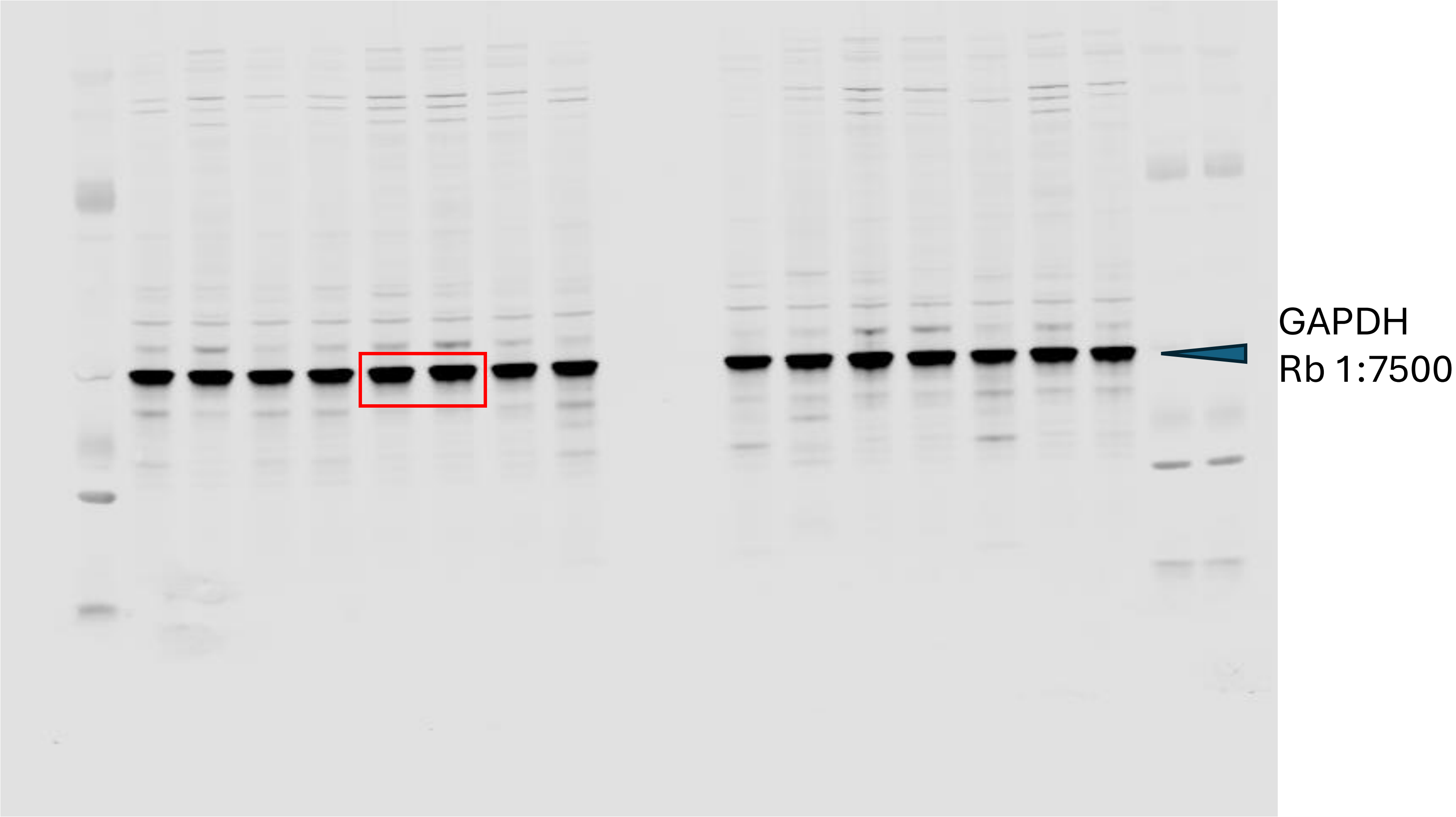


Rb800 ATXN3 & GAPDH (post GAPDH) - brightness & contrast normal to focus on GAPDH
